# RfaH licenses RNA polymerase for long-range transcription

**DOI:** 10.64898/2026.09.18.752654

**Authors:** Luca Buccolieri, Bing Wang, Pim P. B. America, Irina Artsimovitch, David Dulin

## Abstract

Processivity is essential for gene expression: *Escherichia coli* RNA polymerase (RNAP) must transcribe 10+ kbp operons without failure, yet backtrack-induced long-lived pauses threaten premature termination. Gre factors stimulate transcript cleavage to rescue backtracked RNAP, whereas NusA and NusG, which bridge the expressome, respectively stimulate and suppress pausing. How they cooperate to secure full-length transcription is unclear. Using high-throughput magnetic tweezers, we reconstitute *ops*-induced pausing, showing that sequence context sets pause occupancy. RfaH, a NusG paralog recruited at *ops* and essential for long virulence operons, loads onto *ops*-paused RNAP by two pathways: one permitting immediate escape, the other requiring GreA rescue. We find that both NusG and RfaH nullify NusA pause-stimulating effect, RfaH sustaining runs four times longer than NusG and replicating its role *in vivo*, where RfaH depletion leads to conjugation failing. This work provides the foundation for dissecting virulence operon expression and its therapeutic targeting.

## Introduction

Transcription is a fundamental process in gene expression, and its regulation is essential for cellular homeostasis^1–4^. Carried out by RNA polymerase (RNAP), it proceeds through three coordinated stages: initiation, elongation, and termination^5^. Whereas initiation largely determines the level of RNA synthesis, elongation controls its kinetics through DNA-encoded regulatory sequences and transcription factors (TFs)^6–8^. This dynamic phase orchestrates diverse co-transcriptional processes, including translation coupling in Bacteria (**Fig. 1a**) and Archaea, RNA folding of ribozymes and riboswitches and transcription-coupled DNA repair^9–11^. Beyond these roles, elongation dynamics such as nucleotide addition and pauses shape RNAP processivity and promote quality control of nascent transcripts, establishing elongation as a key regulatory step of gene expression^12^.

**Figure 1:**
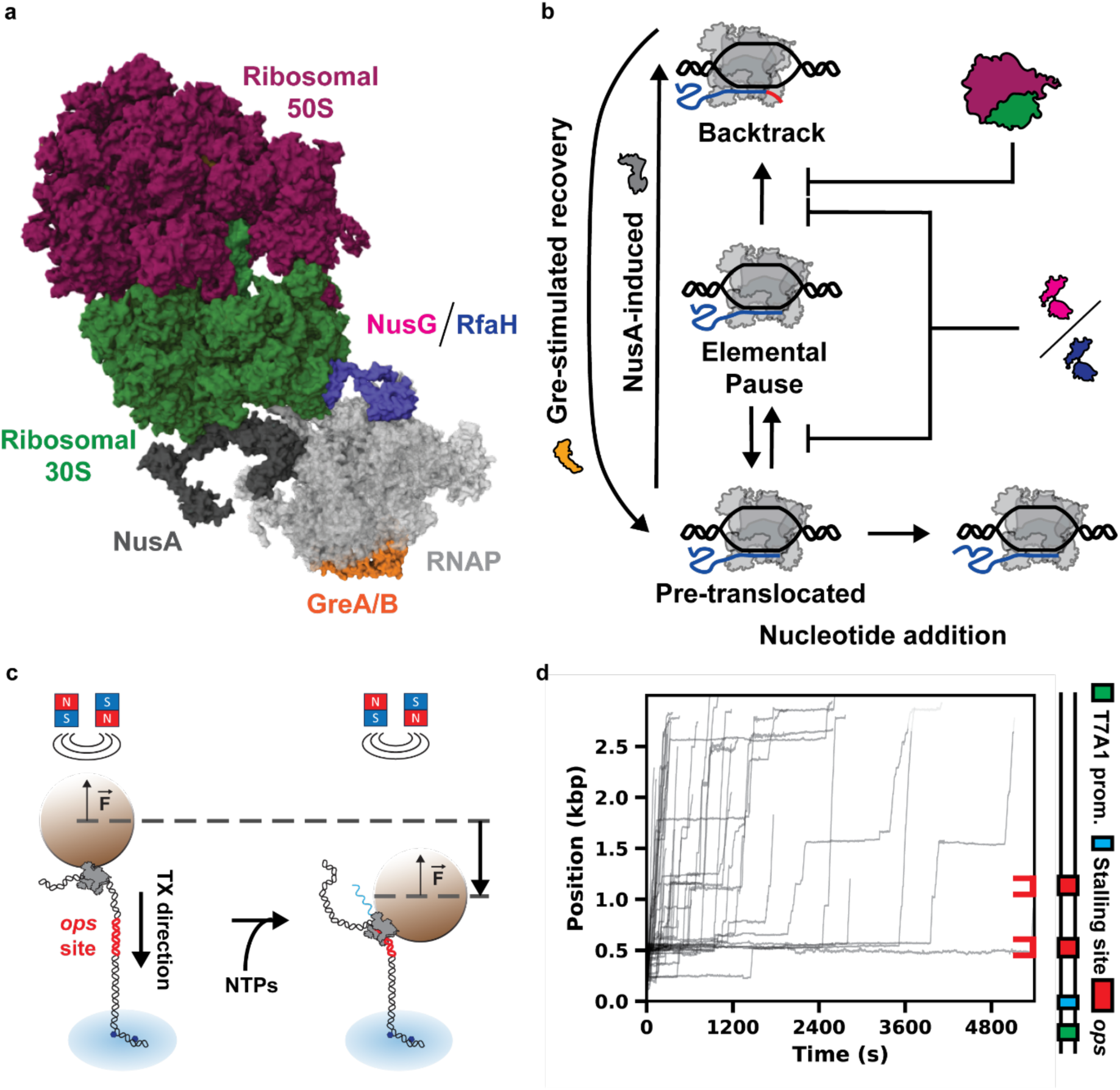
A high-throughput magnetic tweezers assay to investigate the determinants of bacterial transcription processivity. **(a)** Structure of the *E. coli* expressome (PDB entry: 8VOP) showing how RNAP (light gray) is physically coupled to the ribosome small subunit (30S) by the transcription factors NusA (black) and either NusG or RfaH (represented here in violet). Gre (PDB entry: 1GRJ) factors can associate with the RNAP to help recovering from backtracked states. **(b)** Simplified kinetic pathway transcription elongation complex reporting on the different type of pauses interrupting successive nucleotide addition cycles, and the role of the Nus and Gre transcription factors and the ribosome in stimulating or repressing these pauses. **(c)** Schematics of the magnetic tweezers transcription assay for the case the force is opposite to transcription elongation direction. **(d)** Bacterial transcription elongation activity traces obtained under constant opposing force of 7 pN. The DNA template composition is described on the right with a T7A1 promoter (green) and a stalling site (cyan), and two *ops* site (red), at ∼ 0.5 kbp and ∼1.1 kbp downstream the transcription start site.

Pausing has been investigated extensively both *in vivo* and *in vitro*, with pause durations spanning seconds to hundreds of seconds. Single-molecule techniques, including optical, magnetic and nanopore tweezers, have resolved their mechanochemical origin, showing that pauses correspond to off-pathway, catalytically inactive states^13–18^. Three pause classes have been described: the elemental pause, specified by a consensus G_-11_Y_-1_G_+1_ motif^19^ (Y, pyrimidine); the backtrack pause, entered from the elemental state^17^, in which RNAP diffuses backward along the template; and the RNA hairpin- or structure-dependent pauses^20,21^. Single-molecule techniques have shown that RNAP accesses the short-lived, ubiquitous elemental pause from the pre-translocated state of the nucleotide addition cycle, and can either resume elongation or further branch out into a rare, long-lived backtrack state^17,22–25^ (**Fig. 1b**). Among DNA-encoded regulatory pauses, the *ops* element (G_-11_GCGGTAGnnTG_+1_) is the strongest pause signal and serves as a model for this class. When RNAP pauses at the *ops* site, a short hairpin that forms in the non-template DNA strand further stabilizes the pause^26,27^. Distinct *ops* variants are encoded in untranslated leader regions (5’ UTRs) of long virulence and fertility operons^28^, with strictly one copy per operon, and vary in pause occupancy.

Because pauses shape gene expression dynamics, they are tightly controlled by TFs that associate with the elongating RNAP and modulate pause entry and exit rates^29–31^. NusA reduces processivity, either by promoting the elemental pause and thereby increasing backtrack entry, or by stabilizing pause-inducing RNA hairpins and structures^32,33^. Conversely, NusG, the only universally conserved TF, promotes processivity by stabilizing the transcription bubble and preventing RNAP swiveling^34,35^. NusG also recruits Rho termination factor to induce RNA release^36^. RfaH is a specialized paralog of NusG that is exclusively recruited by RNAP transcribing an *ops* sequence^26,37,38^, and is required for the expression of its target operons^37,39,40^. RfaH exerts an anti-pausing effect similar to that of NusG, but binds RNAP with higher affinity and stabilizes the anti-swiveled conformation more durably^35,39,41^. RfaH does not interact with Rho^42^ and instead promotes anti-termination. Furthermore, NusA and either NusG or RfaH bridge the transcribing RNAP to a trailing ribosome (**Fig. 1a**)^11,43–45^. These contacts may facilitate the recruitment of the ribosome to mRNA to initiate translation^46–48^, stabilize the transcription-translation coupled complex throughout protein synthesis, promote processive full-length gene expression by reducing RNAP backtracking^49–52^, and prevent mRNA secondary structures that may impede the trailing ribosome^49^. Gre factors (GreA and GreB) rescue backtracked complexes by stimulating the intrinsic 3’-end transcript cleavage activity of RNAP. They differ in the backtrack depth they target: GreA acts on short backtracks (2–3 bp), whereas GreB rescues deeply backtracked complexes (2–18 bp)^53^. Nus factors and GreA are general TFs virtually represented in all bacterial genomes, are present in the cell at micromolar concentrations^54–57^, and bind to non-overlapping sites on RNAP^34,58^.

Although the effects of individual TFs on RNAP elongation dynamics have been thoroughly characterized by single-molecule approaches^17,41,51,52,59–61^, their coordinated impact on transcription output remains unclear. To address this gap, we used high-throughput magnetic tweezers^17,62,63^ to follow dozens of RNAPs in parallel on kilobase-long templates at millisecond and near base-pair resolution, resolving how Gre and Nus factors and RfaH shape transcription dynamics and processivity with high statistical confidence. We reconstituted strong pausing by the *ops* sequence, with RNAP pauses lasting up to hundreds of seconds. Using two identical *ops* sites separated by ∼0.6 kbp but embedded in different sequence contexts, we found that the first site, in its native context, induces significantly stronger pauses, indicating that sequence context is an important determinant of *ops* pause efficiency. We further found that RfaH associates with *ops*-paused RNAP in two modes: one yielding a pause from which RNAP escapes unaided, and another in which the pause is hyperstabilized by a short backtrack and requires GreA for escape. Most importantly, once released from the *ops* site, the RfaH–RNAP complex counters NusA-induced pausing similarly to NusG, while extending pause-free runs three- to fourfold. Our results indicate that the anti-pausing activity of RfaH favors the expression of long operons by preventing entry into long-lived, backtrack-related pauses. Our study lays the foundation for bottom-up reconstitution *in vitro* of bacterial gene expression in its full complexity.

## Results

### A high-throughput single-molecule assay resolves the *ops*-site pause

We first established a high-throughput magnetic tweezers assay to capture RNAP pausing at the *ops* site and interrogate its mechanochemical origin^17,64^ (**Materials and Methods**, **Fig. 1c**, **Fig. S1a**). In this assay, a bead-bound RNAP stalled in elongation at +29 nt from the transcription start site (TSS) was tethered to the flow-chamber surface through a digoxigenin-decorated DNA handle captured by surface-adsorbed anti-digoxigenin (**Materials and Methods**). The DNA template encodes the *rpoB* gene, interrupted by the *rfaQ* 5′-UTR (the *rfaQ* leader sequence plus the first 200 bp of the gene), with the *ops* site positioned at ∼0.5 kbp, namely *ops1*; an identical *ops* site, here named *ops2*, is located 0.6 kbp downstream (**Materials and Methods**). Depending on whether the handle lies upstream or downstream of the promoter, the force applied to RNAP opposes or assists translocation, respectively (**Fig. S1bc**). In both configurations, dozens of traces were collected per experiment (**Fig. 1d**, **Fig. S1d**), rapidly accumulating large statistics for every condition (**Table S1**) and enabling statistically robust processivity estimation (**Fig. S1e**).

In the activity traces, the *ops1* pause is visually prominent, whereas the pause at *ops2* is far less apparent (**Fig. 1d**, **Fig. S2a-f**). At −7 pN (negative values denote opposing force), the fraction of traces pausing 5 s or more at *ops2* compared to *ops1* is ∼0.5, with median pause durations of 3.28 ± 0.35 s and 4.54 ± 1.39 s, respectively (**Fig. S2a**). This suggests that the transcript sequence itself contributes to pause stabilization, indicating that the *ops* pause depends on the DNA sequence flanking the pause element. The *ops1* pause duration also depends strongly on force magnitude and direction: the median duration decreases almost 3-fold, from 4.54 ± 1.39 to 1.76 ± 0.39 s, as the force is varied from −7 pN to +10 pN (**Fig. S3a, Table S2**), while the RNAP population that escapes the pause increases from 40% to 90% from −10 pN to +10 pN (**Fig. S3b**). This suggests that the *ops* pause is entered from the pre-translocated state (**Fig. 1b**). On the other hand, we did not observe an increase in the escaping population as a function of the force magnitude, indicating that the non-template DNA hairpin at the *ops1* site is not affected by the force.

We observe that Gre factors and NusG prevent RNAP stalling at *ops1* almost entirely (**Fig. S4abc**, **Fig. S5a**), effects likely due to NusG preventing RNAP entry into the backtrack state^41^ and Gre factors rescuing the backtracking RNAP^65^. Conversely, NusA stimulated pausing, doubling the RNAP fraction stalling at *ops1* (**Fig. S4d**, **Fig. S5a**). Furthermore, GreA and NusG do not affect the *ops1* pause duration, while GreB and NusA slightly and significantly increase it, respectively (**Fig. S5b**, **Table S3**).

To probe whether resolving or preventing backtracking recovers the processivity defect induced by NusA, we combined it with either GreA to resolve backtrack, NusG to prevent backtrack, or both GreA and NusG (**Fig. S6a-d**). The two TFs differed strikingly at *ops1* in the presence of NusA: GreA failed to rescue the NusA-induced processivity drop, whereas NusG largely countered it (**Fig. S6e**). However, even the combination of NusG and GreA in presence of NusA is not fully able to recover the drop in processivity at *ops1*. In addition, combining NusA with either NusG, GreA or NusG and GreA does not change the pause duration at *ops1* from the case with NusA alone (**Fig. S6f**, **Table S4**), indicating that NusA dominates the reaction.

In the following sections, we focus exclusively on *ops1*, which provides the clearest read-out of the *ops* pause mechanism.

### RfaH hyperstabilizes backtracked RNAP at *ops* independently of force

We next examined the recruitment and impact of RfaH at the *ops1* site (**Fig. 2a**). RfaH recruitment to RNAP is exquisitely controlled: in free RfaH, the RNAP-binding site on its N-terminal domain (NTD) is masked by the C-terminal KOW domain. When RNAP pauses at *ops*, RfaH NTD binds the non-template DNA hairpin while KOW dissociates and refolds from an α-helical hairpin to a β-barrel, thereby activating RfaH^66^.

**Figure 2:**
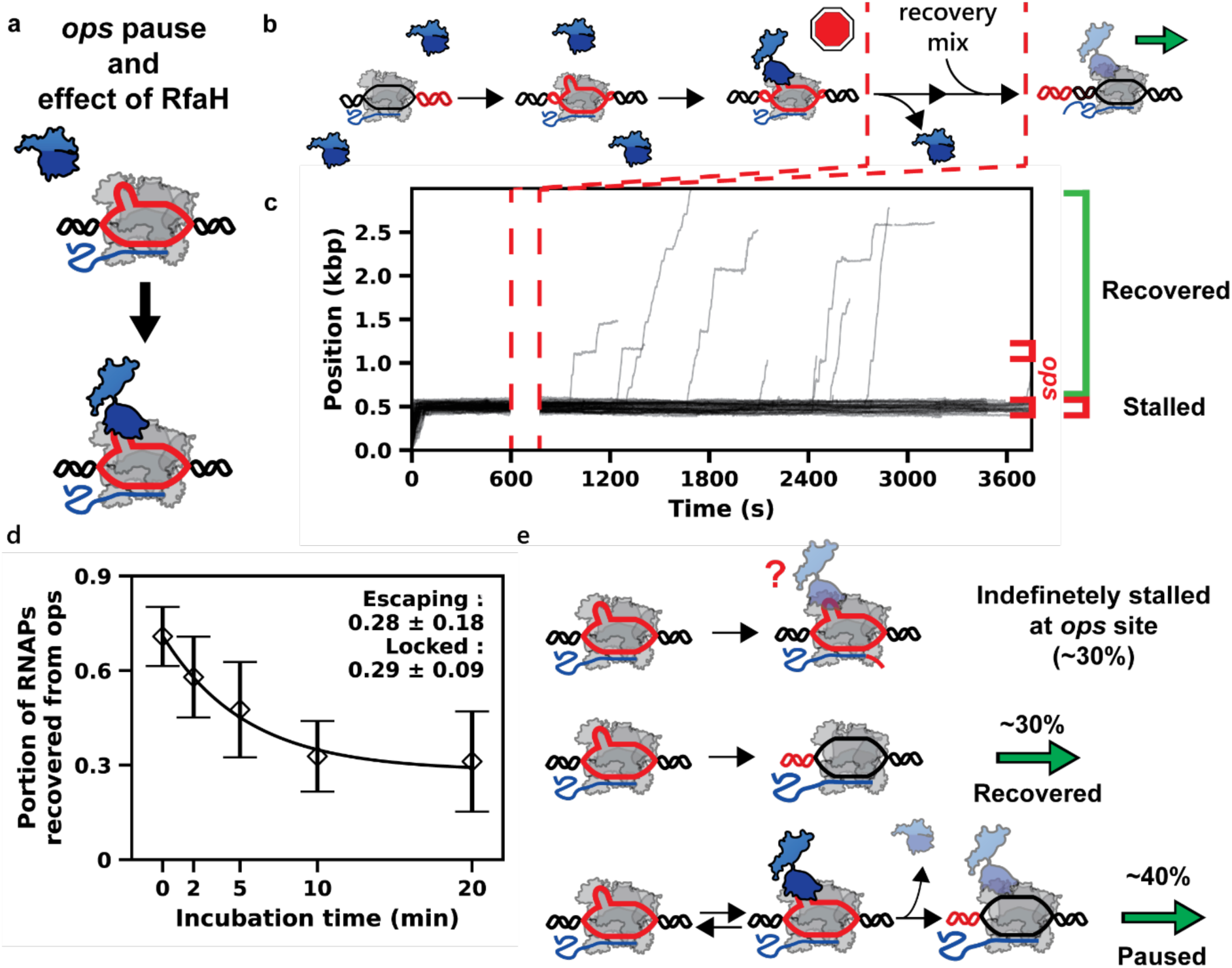
RfaH hyperstabilizes RNAP at the *ops* site in a time-dependent fashion. **(a)** Representation of the RNAP pausing at *ops* to recruit RfaH (KOW domain in light blue, NGN domain in dark blue). **(b)** Schematics of the single-molecule recovery assay: RNAP is incubated with 1 µM RfaH, followed by rinsing and restart of activity with 1 mM NTP and **(c)** exemplary activity traces. The gap indicates rinsing and supplementation of the reaction mix. **(d)** Scatter plot comparing the populations escaping after 2-20 min incubation. The error bars represent the Wilson score with 95% CI. **(e)** Model describing the behavior of RNAP at *ops* with saturating RfaH, defining 2 fixed populations, one that stalls indefinitely and one that always recovers from the *ops* site, and a third one (paused) sensitive to incubation time.

RfaH visibly increased the number of traces stalling at *ops1* (**Fig. S7a-d**). In an RfaH titration, the fraction of RNAPs escaping the *ops1* pause decreased above 5 nM (**Fig. S8a**), consistent with a nanomolar binding affinity. However, there was no significant change in the median pause duration at *ops1* (**Fig. S8b**, **Table S5**). We then assessed the force dependence of *ops1* escape at saturating RfaH (1 µM), varying the force from −10 to +10 pN (**Fig. S9a-e**). Surprisingly, the fraction of RNAPs escaping *ops1* was essentially force independent, i.e. (54±12)% (mean±s.d.) (**Fig. S9f**), though ∼25% lower than without RfaH (**Fig. S9g**). Furthermore, the median *ops1* pause duration was also unaffected by force (**Fig. S10a**, **Table S6**). Although assisting force should disfavor the pre-translocated state required to enter this pause, hyperstabilization persisted, indicating that RfaH recruitment is governed primarily by molecular recognition. It also implies that force plays at most a minor role in *ops1* escape, within the range tested and at saturating RfaH.

To test whether hyperstabilization depends on incubation time and on the continued presence of RfaH in solution, we performed a two-step recovery assay in which RfaH was removed after 2–20 min of incubation (**Fig. 2bc**). Three subpopulations emerged: ∼30% of RNAPs escaped *ops1* regardless of RfaH, ∼30% never escaped under any condition, and the remaining fraction escaped in a time-dependent manner (**Fig. 2d**), spreading escape across the experimental time scale, unlike the case in which RfaH was absent from solution (**Fig. S11a**). Re-injecting RfaH after a short incubation and rinse recapitulated the long-incubation phenotype (**Fig. S11b**), indicating that RNAP escape follows RfaH dissociation. Together, these results establish that RfaH binds RNAP at *ops1* with high affinity, irrespective of force direction and magnitude, and that prolonged co-incubation or re-exposure progressively drives a conformational change that locks RNAP into a hyperstabilized state (**Fig. 2e**), from which escape requires the intervention of other factors. If RfaH dissociates before hyperstabilization, the KOW will refold and rebind NTD, reestablishing the autoinhibited state^67^.

### GreA as the main driver of escape from the RfaH-hyperstabilized *ops* pause

Next, we asked what mechanism allows RNAP to productively escape the hyperstabilized pause state (**Fig. 3a**). The RfaH–hairpin interaction drives the elongating RNAP into a short backtrack^66^; any reaction that restores the complex to an elongation-competent state should therefore promote productive escape. We therefore tested the effect of Gre factors or of a trailing, non-biotinylated RNAP (nbRNAP) on the stalled complex (**Fig. 3a**, **Fig. S12abc**). In both cases, the fraction of RNAPs transcribing beyond *ops1* increased substantially, from ∼55% to ∼80% (**Fig. 3b**). However, ∼20% of RNAPs failed to recover from the *ops1* pause in the presence of Gre factors (**Fig. 3b**), whereas most complexes escaped in the absence of RfaH (**Fig. S5a**), suggesting a state inaccessible to Gre-stimulated cleavage.

**Figure 3:**
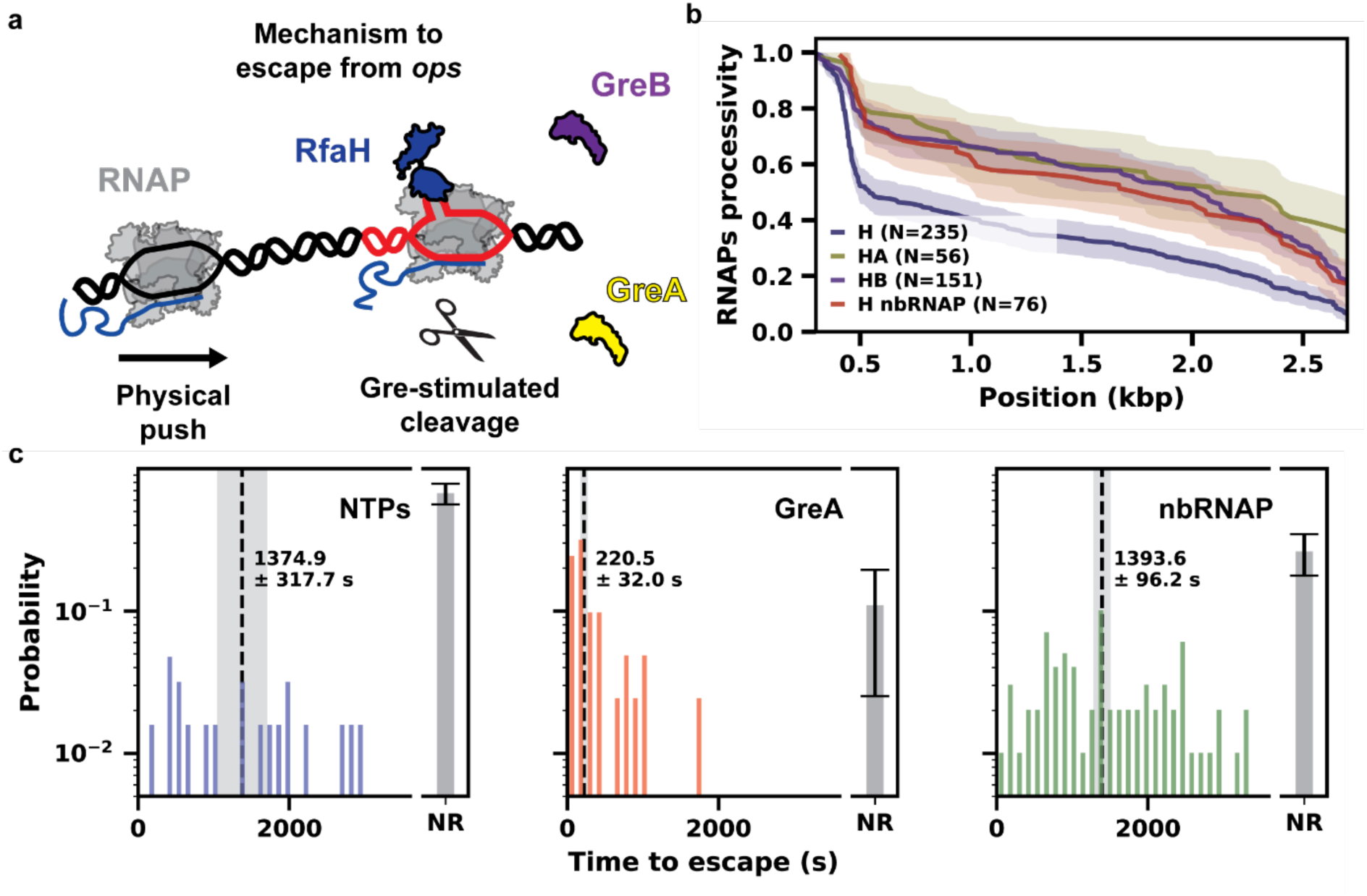
Productive escape of the RfaH-RNAP complex from the *ops* site is prompted by GreA. **(a)** Representation of the different mechanisms RNAP uses to escape from *ops* after loading RfaH: a physical push, i.e non-biotinylated RNAP (nbRNAP), or Gre-stimulated RNAP intrinsic RNA cleavage. **(b)** RNAP processivity with 1 µM RfaH (H) in isolation or with either 2 µM GreA (A) or 2 µM GreB (B) or 10 nM nbRNAP. **(c)** Histogram (2 min bins) of time to escape from *ops* during recovery assay after 10 min RfaH incubation), resuming transcription with either NTP only, with 2 µM GreA, or 10 nM nbRNAP. The median of the observed durations is indicated with a dashed vertical black line, with the light grey range indicating one standard deviation of the medians obtained from 20 bootstrap procedures. The grey bars represent the stalled population, and the error bars are the Wilson score with 95% CI.

Surprisingly, the median *ops1* pause duration increased 30-fold when RfaH was combined with GreA, and only mildly — though significantly — with GreB or nbRNAP (**Fig. S12d**, **Table S7**). This suggests that GreA rescues the RfaH–RNAP paused complex by a mechanism distinct from that of GreB or a trailing nbRNAP, requiring the complex to equilibrate into a state in which the transcript becomes accessible to GreA-stimulated cleavage. In the recovery assay (**Fig. 3b**), after 10 min of RfaH incubation, GreA returned ∼90% of complexes to elongation, compared with ∼70% for a trailing nbRNAP and ∼30% for NTPs alone (**Fig. 3d**). The latter two spread escape over longer and distinct time scales, whereas GreA confined it to within 10 min, implying that hyperstabilized complexes eventually settle into a short backtrack readily rescued by GreA.

In addition, as NusA impede the RNAP transcribing the *ops1* site by inducing stalling, we asked how the presence of RfaH would influence the RNAP behavior. At low RfaH (5 nM) and saturating NusA (**Fig. S13a**), RfaH prevented the drop in processivity at *ops1* seen with NusA alone (**Fig. S13b**), the latter dominating the median *ops1* pause duration (**Fig. S13c**, **Table S8**). This suggests that transient RfaH binding at *ops1* prevents entry into the deep backtracked state stimulated by NusA, lowering the barrier for RNAP to return to a shallow backtrack. Adding NusA and GreA together (**Fig. S14a**) significantly affected neither the stalling at the *ops1* site (**Fig. S14b**) nor the median pause duration of the hyperstabilized RfaH– RNAP complex (**Fig. S14c**, **Table S9**). In conclusion, our data are consistent with RfaH first forming an unstable, reversible complex with the *ops1*-paused RNAP, which either converts into a hyperstabilized state requiring GreA for escape, or escapes directly and loses RfaH shortly thereafter. The remaining stalled RNAPs constitute an unrecoverable population.

### Gre factors with either NusG or RfaH reverse the NusA-induced processivity loss

Having established the conditions for productive escape from *ops1*, we next investigated RNAP processivity and elongation dynamics downstream of the pause. As a benchmark, we first measured the processivity of RNAP alone as a function of force (**Fig. S3c**). Downstream of *ops1*, however, opposing forces below 10 pN did not impair processivity (**Fig. S3c**). To analyze elongation dynamics quantitatively, we used a dwell-time analysis, in which the distribution of times to incorporate ten consecutive nucleotides (the dwell time) is obtained by scanning each trace with a non-overlapping 10 nt window (**Fig. S1fg**). We fitted the distribution with a stochastic pause model comprising a gamma distribution for pause-free nucleotide addition, two exponential distributions for the short (∼0.5 s) and medium (∼2.5 s) pauses, termed Pause 1 and Pause 2, respectively, and a power-law distribution with a −−3/2 exponent for the long-lived, backtrack-related pauses^62,68^. The power-law probability was the most force-sensitive parameter (**Fig. S3d-j**), decreasing from ∼0.01 to ∼0.002 as the force was varied from −10 to +10 pN, consistent with previous reports^69^.

We next evaluated the individual impact of NusA, NusG, RfaH, GreA and GreB at physiological concentrations on processivity and elongation dynamics. NusA consistently reduced RNAP processivity (**Fig. S5c**). This effect is apparent in the activity traces (**Fig. S4d**) and arises from an increased probability of pause entry (**Fig. S5hi**) and, consequently, a higher backtrack probability (**Fig. S5j**). NusG, like NusA, decreased the pause exit rates (**Fig. S5fg**) but increased only the Pause 1 probability (**Fig. S5h**). RfaH did not significantly alter RNAP elongation kinetics at any concentration (**Fig. S8d-j**) or force (**Fig. S10b-h**) tested. GreA reduced the probabilities of Pause 2 and long-lived pauses by ∼30% (**Fig. S5ij**), whereas GreB slightly inhibited elongation (**Fig. S5e-h**), as previously reported^59,70,71^, and lowered the likelihood of long-lived pauses by 60% (**Fig. S5j**). Under opposing force, the dominant bottleneck for processivity is entry into deep backtracked states that cannot be recovered, even by RNA cleavage, probably because the RNA 3’-end is no longer positioned in the catalytic site or has been dislodged into the secondary channel^53,72^. In conclusion, none of these TFs improved RNAP processivity individually.

For RNAPs that overcame the *ops1* pause in the presence of a combination of TFs (**Fig. 4a**), adding either NusG or GreA improved processivity in the presence of NusA, with an additive effect when NusG and GreA are combined (**Fig. 4bc**). The dwell-time analysis, however, shows that NusA still increases the probability of pause entry and the pause duration even with NusG and GreA present, partially decoupling pause dynamics from processivity (**Fig. S6g-m**). Opposingly to NusG, which can reassociate after dissociation from the RNAP, adding RfaH in the presence of NusA showed a faster decay in processivity after 1 kbp (**Fig. 13d**), consistent with the inability of RfaH to load again on the RNAP. Accordingly, the elongation dynamics are impaired by the pausing effect of NusA (**Fig. 13e-k**).

**Figure 4:**
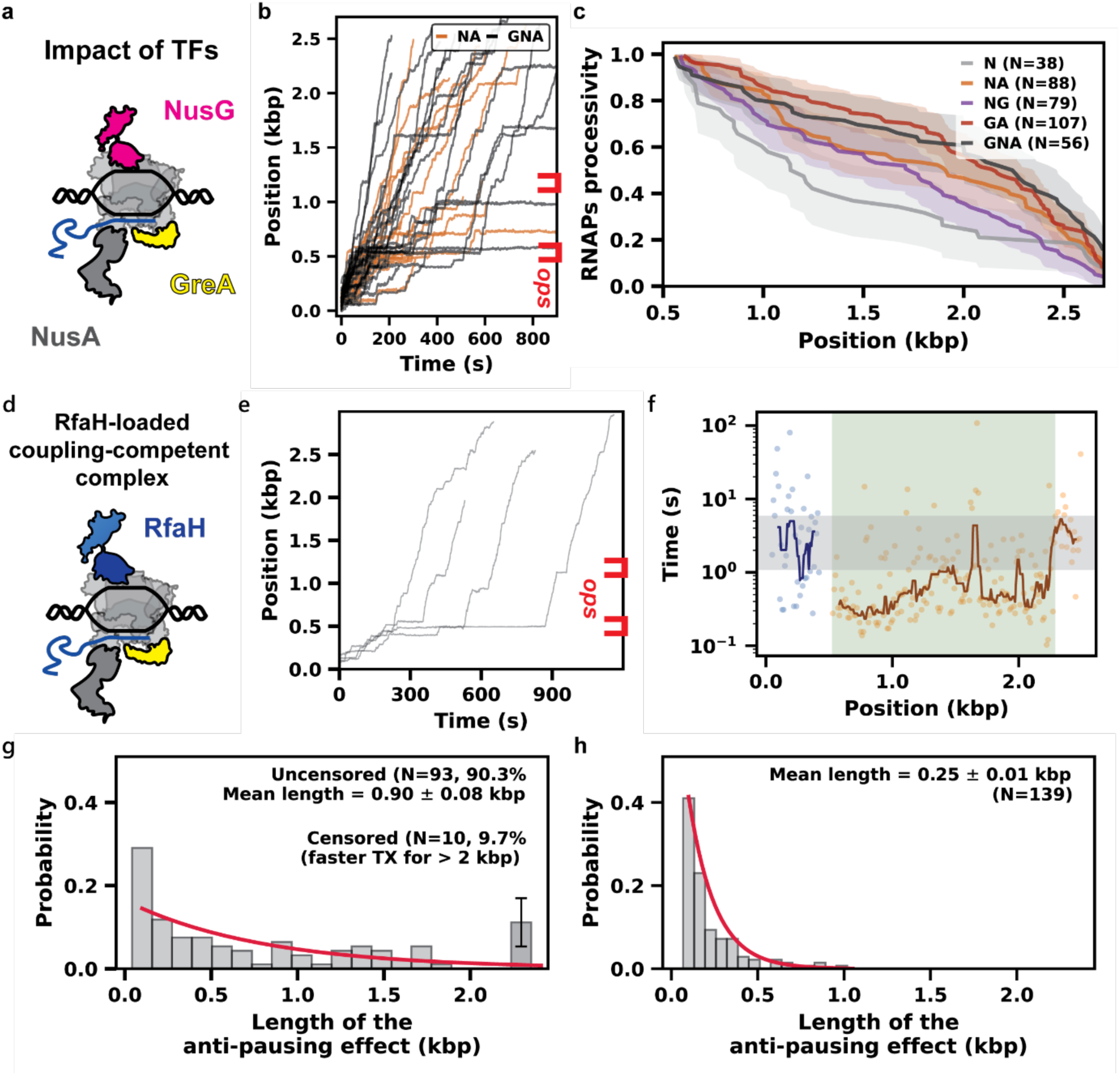
RfaH replaces NusG to counter NusA deleterious effect on RNAP processivity after GreA-stimulated escape from the *ops* site. (a) Schematic representation of the RNAP transcribing in the presence of GreA (A), GreB (B), NusA (N) and NusG (G). (b) Comparison of 30 exemplary traces in the presence of NusA and GreA in the absence (NA) or presence of NusG (GNA). (c) Processivity of the RNAP for 7 pN OF after the *ops* site with NusA (N) in combination with either GreA (NA) or NusG (GA), the combination of the latter two (GA) or the three TFs present together (GNA). (d) Schematic representation of elongating RNAP associated with RfaH and NusA. (e) 4 traces with RfaH, GreA and NusA, *ops* site in red. (f) Dwell times (dots) as a function of the RNAP position along the template for a single trace, before (blue) and after (orange) the *ops* site. Trace collected in the presence of RfaH, NusA and GreA. The solid line is a 10-point rolling median. The grey shaded area is the 95% CI of the medians from 2000 bootstrap procedures extracted from the dwell times preceding the *ops* site (∼0.5 kbp). The green shaded area is the region of the trace where the median does not cross the the lower end of the CI for more than 10 dwell times. (g) Histogram of the pause-free distance as defined for the green shaded area in (f), using 2 µM GreA, 200 nM NusA and either (f) 1 µM RfaH or **(h)** 1 µM NusG. The black bar in (f) represent the fraction of traces where the pause-free area continued until the end of the template (censored).

We then asked how RfaH compares to NusG in a reaction with NusA and GreA (**Fig. 4d**). We found that processivity was similarly impaired with either RfaH or NusG when NusA was present (**Fig. S14d**), probably because RfaH dissociates. Consistent with the *ops* specificity of RfaH, the elongation dynamics changed downstream of *ops1* (**Fig. 4e**), with a decrease in the median time to transcribe 100 bp (**Fig. 4f**) followed by an eventual switch back to a pause-prone state, indicating that RfaH had dissociated. Eighty percent of RNAPs were affected by RfaH (**Fig. S15a**), transcribing faster from *ops1* until RfaH dissociation. RfaH loading conferred a clear kinetic advantage relative to the region upstream of *ops1* (**Fig. S15b**), reproducing the effect of NusG. Relative to the region upstream of *ops1*, segments with a decreased median transcription time showed a slightly faster nucleotide addition rate (**Fig. S15c**) and Pause 1 exit rate (**Fig. S15d**), together with a 3-fold reduction in the probabilities of Pause 2 and of long-lived pauses (**Fig. S15fg**). These results support a model in which RfaH and NusG similarly prevent pauses from limiting processivity.

We next compared the distance covered during anti-pausing events with RfaH vs NusG. Although the elongation dynamics were comparable between the two TFs, RfaH increased the run length free of long-lived pauses 3- to 4-fold (0.90 ± 0.08 kbp, **Fig. 4g**) relative to NusG (0.25 ± 0.01 kbp, **Fig. 4h**). Notably, ∼10% of the population with RfaH transcribed without long-lived pauses until the end of the template, showing particularly strong association with the transcribing RNAP.

We finally probed whether our observations are robust *in vivo*, where the translating ribosome is expected to further stabilize the RfaH-RNAP complex by sequestering the KOW domain. We explored RfaH regulation of conjugation of F, a single-copy 100-kb IncF plasmid that carries the 32-kb *tra* operon, the longest in *E. coli*, transcribed from the P_Y_ promoter.

Transcription from P_Y_ is activated by the F-encoded TraJ and host co-activator ArcA^73^ and is repressed by H-NS^74^. The post-initiation control of the *tra* operon, which has an *ops* element downstream from the *traV* gene (**Fig. 5**) has not been investigated, despite its massive length and an early report that *tra* expression depends on RfaH^75^.

**Figure 5:**
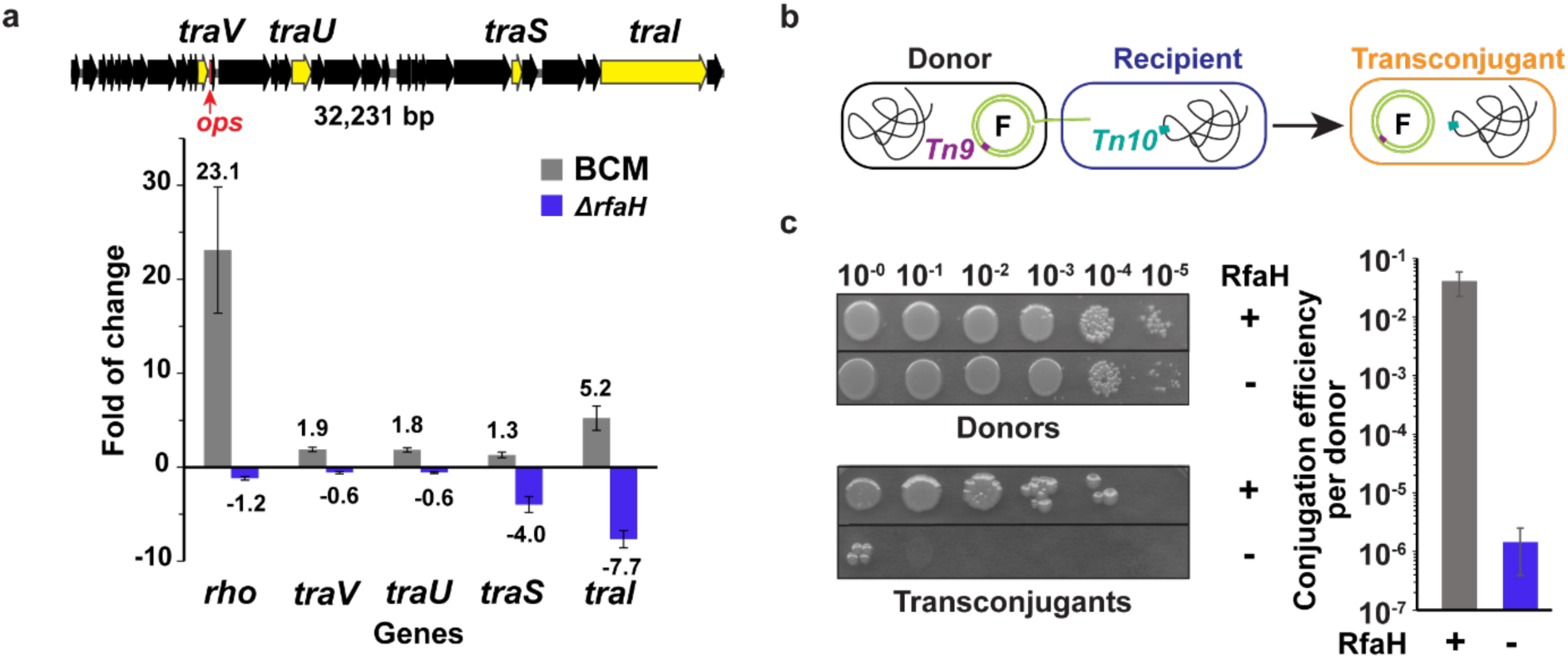
Effects of RfaH on the F-factor *tra* operon. **(a)** Effects of Rho and RfaH on the F-factor *tra* operon expression were tested using RT-qPCR. Fold of change is calculated relative to the wild-type *E. coli* strain. BCM, bicyclomycin. Mean fold changes are shown above the bars. Negative values represent reduced expression. Error bars represent the SD of three biological replicates. **(b)** Schematic of conjugation assay. **(c)** Representative plates from four independent biological replicates are shown. Serial dilutions are indicated. Conjugation efficiency per donor is plotted on a log10 scale. Error bars represent SD (n = 4).

We hypothesized that, similarly to other long *ops*-containing *E. coli* operons^42,76,77^, the F *tra* operon is silenced by Rho and counter-silenced by RfaH. Indeed, we found that the deletion of *rfaH* reduced the expression of distal *traS* and *traI* genes by 4- and 8-fold (**Fig. 5a**). The observed polarity was reversed in the presence of bicyclomycin (BCM), a specific inhibitor of Rho; the *traI* expression was increased 5-fold when Rho was inhibited by BCM (**Fig. 5a**). A mating assay shows that the deletion of *rfaH* causes a dramatic, ∼28,000 fold decrease in conjugation efficiency (**Fig. 5bc**), suggesting that RfaH activation of translation makes the decisive contribution to its counter-silencing activity. This result is consistent with our previous findings^48^ and suggests that RfaH loads the initiating ribosome and couples it to the transcribing RNAP over long distances whereas NusG, which might form a complex with RNAP when RfaH is absent, is unable to substitute for RfaH. The *ops* element is present in all F-like plasmids^78^, suggesting that this mode of regulation is widespread.

Overall, these observations indicate that the anti-backtracking activity of RfaH counteracts the pause-inducing behavior of NusA *in vitro* and the silencing exerted by H-NS *in vivo*, and that its stable association with RNAP at *ops* site and downstream makes it more robust in transcribing long operons than NusG, which exchanges frequently (**Fig. 6**).

**Figure 6:**
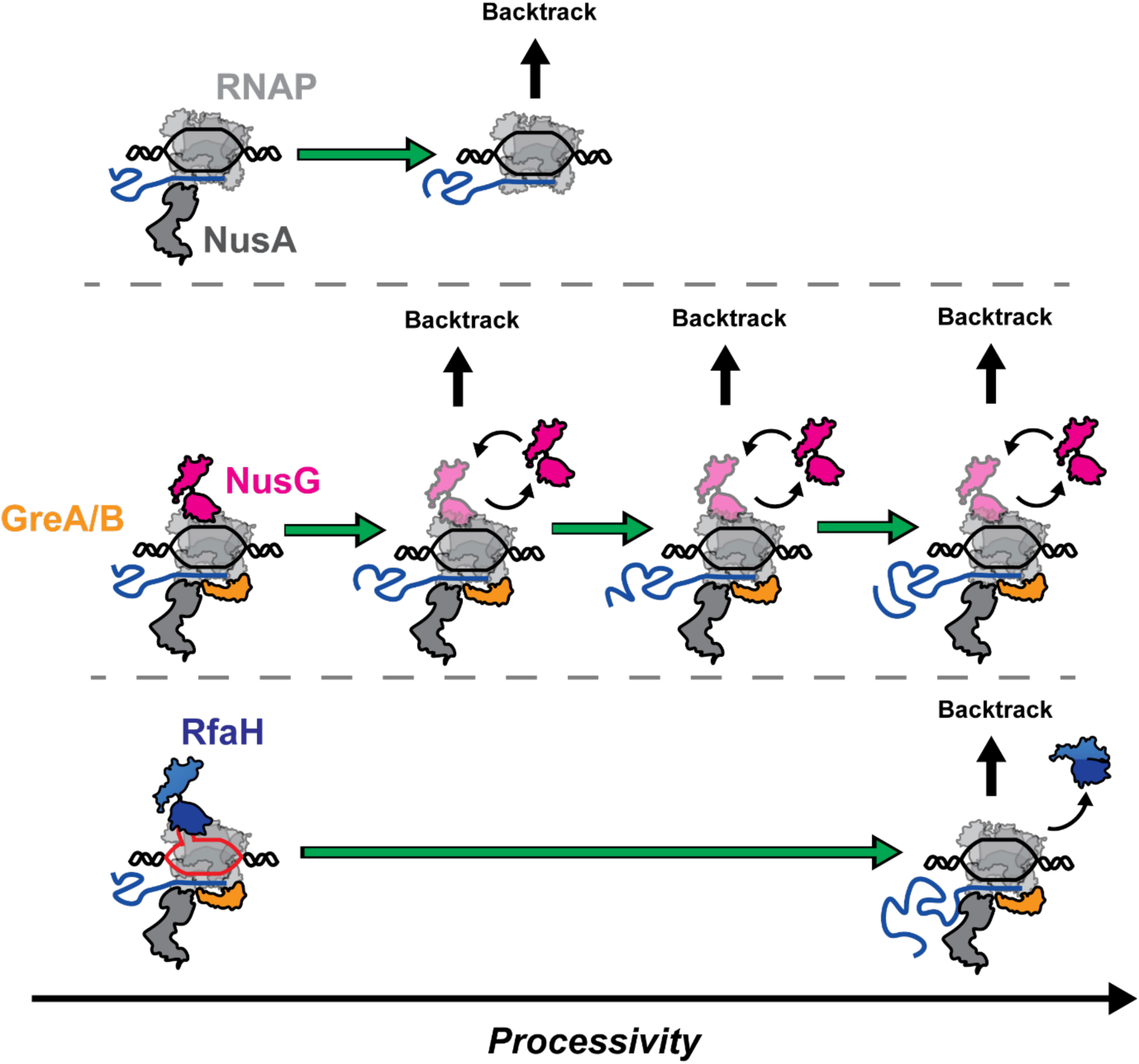
RNAP processivity is augmented by the assembly of TFs that preserve regulatory pauses and limit deleterious backtracking. The RNAP in the presence of NusA regularly pauses, eventually resulting in premature termination through entering a backtrack state (top). The addition of NusG partially restore the processivity by limiting the entry NusA-stimulated pauses, but the frequent exchange of NusG leads to regular entry into long-lived, backtrack pauses (middle). On the other hand, the stable association of RfaH with RNAP strongly reduces the entry into pauses, assisting long transcript synthesis (bottom).

## Discussion

Here we established a high-throughput magnetic tweezers assay to dissect the *ops* pause and its role in recruiting RfaH, a NusG paralog essential for the expression of virulence operons in several Gram-negative pathogens. We then asked how Nus and Gre factors affect the *ops* pause in the presence and absence of RfaH, and how they jointly shape the dynamics and processivity of transcription (**Fig. 1**). We found that the *ops* pause is sequence-context dependent, with a mean duration of ∼100 s at 7 pN opposing force. NusA further populates this pause, whereas NusG and GreA allow RNAP to escape rapidly. RfaH associates with *ops1*-paused RNAP in two successive modes: an initial reversible complex and a subsequent hyperstabilized state (**Fig. 2**). Escape from the latter requires GreA (**Fig. 3**). The pause-inducing activity of NusA reduces RNAP processivity, an effect partially countered by either NusG or RfaH together with GreA (**Fig. 4**). Finally, the stable association and anti-pausing activity of RfaH sustain longer runs free of long-lived, backtrack-related pauses, favoring transcription of long operons *in vivo* (**Fig. 5).**

Previous single-molecule work reported a much shorter, unstable *ops* pause, with a mean duration of ∼4 s at 7 pN assisting force ^79^. Our data show a mean *ops1* pause duration of ∼15 s at the same assisting force, a nearly 4-fold difference. Two differences may account for this. First, the *ops* site used by Herbert and colleague comes from a different operon (*pheP* vs. *rfaQ*) and was inserted directly into *rpoB* gene (used as template) without its native flanking region; natural *ops* variants differ in strength, so sequence alone could contribute. Second, we show that an *ops* site lacking its native flanking region gives a shorter pause (*ops2* vs. *ops1*), which could further account for the discrepancy. How the *rfaQ* leader sequence stabilizes the *ops* pause remains unclear. It is possible that DNA sequences or RNA secondary structures further stabilize the *ops* pause, e.g. by stimulating entry into a backtrack state^80^.

Recent structural and biochemical work showed that RfaH binding to RNAP can drive the complex into a backtracked state that requires GreA for escape^66^, consistent with our observations. That study also reported that NusA further stabilizes the RfaH-mediated pause, again in agreement with our data. However, those complexes were pre-assembled in solution for cryo-EM imaging rather than formed by RNAP transcribing through *ops*. Our measurements on actively transcribing complexes resolve at least three distinct RfaH–RNAP species: an unstable complex, a hyperstabilized pause that requires GreA for escape, and a terminally stalled state at *ops1*. Formation of the hyperstabilized complex is slow and yields the most stable and processive RNAP, as reflected in its counteraction of the NusA pause-stimulating activity.

Although NusG and RfaH belong to the same superfamily and both exert anti-pausing effects on RNAP, they differ in two key respects: RfaH makes stronger contacts with RNAP than NusG does, and NusG can recruit Rho to promote termination whereas RfaH cannot. RfaH is also present at tens of nanomolar *in vivo*, far below the micromolar concentration conventionally assigned to NusG. However, only ∼10% of cellular NusG is free, the remainder being bound to molecular machines or to DNA^81^. This both limits the pool of NusG available for any given complex (termination or anti-termination) and keeps a fraction pre-positioned near the transcription machinery. Similarly, NusA is present at a few hundred nanomolar^57^, whereas RNAP reaches single-digit micromolar concentrations, of which 20-30% are actively transcribing at any time^82^, suggesting that the available NusA fraction is limiting *in vivo*. Our experiments at saturating NusA therefore represent an extreme case unlikely to occur in the cell, and the processivity we measure with NusA is most likely a lower bound.

RfaH is required for expression of long horizontally transferred operons that lack canonical ribosome binding sites and are silenced by H-NS^77^. RfaH-KOW binds ribosome to activate translation^83^ and counter-silences H-NS^76,77^. Our results reveal a counterintuitive but coherent role for NusA within RfaH-regulated operons. On its own, NusA is a potent inhibitor of processivity, driving RNAP into pause states that may serve as intermediates for backtracking. In RfaH-regulated operons, NusA would be expected to compound the silencing effect of H-NS on elongation. Instead, RfaH largely suppresses NusA-induced pausing, attenuating its cost to processivity while preserving NusA-dependent bridging of the ribosome to RNAP^84^. Expressome-mediated gene expression has recently been demonstrated at the single-molecule level for NusA- and NusG-bridged complexes^52^, but not for RfaH. Because RfaH bridges the ribosome and RNAP more tightly than NusG, it may sustain coupling over longer distances, and the bridge would itself stabilize RfaH on RNAP by preventing KOW fold switch; consistently, RfaH remains bound to RNAP transcribing the *rfa* (*waa*) and other operons *in vivo*^85^. Future work will aim to reconstitute an RfaH-bridged expressome and to determine how such a complex sustains expression of 10+ kb operons.

Crucially, our *in vivo* observations (**Fig. 5**) define the possibility of a general mechanism of regulation of F-like plasmids, all encoding a single *ops* site, through RfaH-mediated boost in RNAP and ribosome processivity. We show that RfaH is critical for plasmid conjugation, with the four orders of magnitude decrease when RfaH is absent. Plasmid conjugation is the primary mechanism of gene transfer in bacterial populations, and epidemic IncF plasmids commonly encode several antimicrobial resistance and virulence determinants^86^.

Studies of conjugation control may expose vulnerabilities that can be leveraged to combat the existential threat of antimicrobial resistance.

In addition to its importance for antimicrobial resistance, RfaH could serve as a model peptide for synthetic biology and for the construction of a synthetic cell^86^. Indeed, RfaH could help overcome the bottleneck imposed by the low efficiency of protein synthesis in vitro compared to *in vivo* expression levels. Although previous efforts have achieved higher yields^87^, translation processivity remains strongly impaired^88^, resulting in truncated proteins and inefficient substrate use. Reconstituting an RfaH-RNAP complex in a synthetic cell would offer two advantages: full-length transcription and translation of long synthetic operons, and the extremely low factor concentration required to achieve it.

Several limitations constrain the scope of our conclusions. On the biological side, while the opposing-force regime emulates unfavorable conditions for transcription, it may not fully recapitulate the mechanism by which RfaH–RNAP complexes transcribe through H-NS filaments *in vivo*. On the experimental side, RfaH loading and its downstream effects were inferred indirectly from RNAP dynamics, without a direct readout of factor occupancy. This prevents us from quantifying the length of template over which RfaH remains bound to RNAP, and from attributing the counteraction of NusA specifically to the anti-backtracking activity of RfaH, to competition with NusA for the β flap tip, or to both. Finally, our reconstitution omits three key *in vivo* players: H-NS, responsible for silencing of the virulence genes, the ribosome, essential for transcription–translation coupling, and the termination factor ρ, which would expose a further functional difference between NusG and RfaH. Direct read-out from fluorescently labeled factors will be needed to resolve these contributions and to clarify how RfaH enhances transcription and translation *in vivo*.

## Materials and Methods

### Expression and purification of *E. coli* RNAP and transcription factors

#### *E. coli* biotinylated RNAP core

Plasmid encoding core RNAP [pIA1202; α-β-β’[AVI][His]- ω] under control of T7 promoter were transformed into *E. coli* BL21(λDE3) and grown in Terrific Broth at 37°C to OD_600_ ∼0.5. Protein expression was induced with 0.5 mM IPTG for 5 hours at 30°C. *E. coli* biotin ligase BirA (Addgene#109424) in *E. coli* BL21(λDE3) was expressed using the same induction condition. Cells were pelleted at 6,000 x g, 4°C for 10 min. To increase the fraction of biotinylated RNAP, cell pellets containing RNAP core and BirA were mixed and resuspended in Lysis Buffer (10 mM Tris-OAc pH 7.8, 0.1 M NaCl, 10 mM ATP, 10 mM MgOAc, 100 μM d-biotin, and 5 mM β-ME) supplied with Complete EDTA-free Protease Inhibitors (Roche) per manufacturer’s instructions.

Cells were opened by sonication, and cell debris was pelleted by centrifugation (20,000 x g, 40 min, 4°C). The cleared cell lysate was incubated with Ni Sepharose 6 Fast Flow resin (Cytiva) for 40 min at 4°C with agitation. The resin was washed stepwise with Ni-A Buffer (25 mM Tris-HCl, pH 6.9, 5% glycerol, 500 mM NaCl, 5 mM β-ME, 0.1 mM phenylmethylsulfonyl fluoride (PMSF)) supplied with 10 mM, 20 mM, and 30 mM imidazole. Protein was eluted in Ni-B Buffer (25 mM Tris-HCl, pH 6.9, 5% glycerol, 5 mM β-ME, 100 mM NaCl, 300 mM imidazole). The eluted protein was diluted 1.5 times with Hep-A Buffer (25 mM Tris-HCl, pH 6.9, 5% glycerol, 5 mM β-ME) and then loaded onto Heparin HP column (Cytiva). A linear gradient between Hep-A and Hep-B Buffer (25 mM Tris-HCl, pH 6.9, 5% glycerol, 5 mM β-ME, 1 M NaCl) was applied. The biotinylated RNAP core is eluted at ∼40 mS/cm. The eluate from Heparin HP column was diluted 2.5 times with Hep-A Buffer and loaded onto Resource Q column (Cytiva). A linear gradient was applied from 5% − 100% Hep-B Buffer. The biotinylated RNAP core was eluted at ∼25 mS/cm. Fractions from the elution peaks were analyzed by SDS-PAGE. Those containing purified protein were combined and dialyzed against Storage Buffer (20 mM Tris-HCl, pH 7.5, 150 mM NaCl, 45% glycerol, 5 mM β-ME, 0.2 mM EDTA).

The σ70 (pIA1127) was purified following previously published protocol^90^. RNAP holoenzymes were assembled by mixing the core RNAP with a 3-fold molar excess of σ^70^, followed by incubation at 37°C for 15 min.

NusG (pIA247) was prepared following^91^. RfaH (pIA238) and NusA (pIA370) were purified following the previous protocol^66^. GreA (pIA578) and GreB (pIA576) were prepared as before^92^.

#### Reverse transcription quantitative PCR (RT-qPCR)

*E. coli* MG1655 derivatives carrying F-factor (IA1016, IA1017; **Table S12**) were cultured in LB with chloramphenicol. The cells were grown at 37 °C to an OD600 of ∼0.7 and treated with 50 μg/mL bicyclomycin (BCM) for 15 min when indicated. The cells were mixed with 0.2 volume of ice-cold growth stop solution (5% phenol and 95% ethanol) to inactivate cellular RNase before being collected by centrifugation at 4,000 × g for 10 min. The mRNA was isolated using the Monarch Total RNA Miniprep Kit (NEB, cat# T2010S). RNA samples were treated with TURBO DNase (ThermoFisher, cat# AM2239) before RT-qPCR. A total of 100 ng RNA samples were used with the iTaq Universal SYBR Green One-Step Kit (Bio-Rad, cat#1725150) and analyzed on a CFX96 system (Bio-Rad). Samples without reverse transcriptase were used as a negative control to ensure the absence of DNA contamination. The quantification cycle (Cq) values were calculated using CFX Manager v3.0 in regression mode. The gene expression level was analyzed by the threshold cycle (2−ΔΔCT) method (PMID: 11846609); the *ihfB* gene was used as a reference, expression of which is not affected by BCM (PMID: 33460557).

#### Conjugation assay

*E. coli* MG1655 strains harboring the F factor were used as donors, and the *E. coli* MG1655 *argE*::Tn10 strain was used as the recipient. Donors and recipients were grown in LB supplemented with selective antibiotics at 37 °C, 250 rpm. Cells were collected by centrifugation when OD reached 0.8 – 1.0, and the pellet was washed once with sterile PBS. Following the wash, the donor and recipient cells were mixed to get ODs of 20 and 40, respectively. 20 μl of mixture was dropped onto an MCE membrane filter (0.22 μm pore size; Millipore Sigma, cat# GSWP01300) overlaid on LB agar and incubated at 30 °C for 40 min (*rfaH*+ strain) or 2 h (*rfaH*-strain). The filter paper was then vortexed in 1 ml of sterile PBS. Serial dilutions of each sample were spotted on M9 (ATCC Medium 2511) and LB + chloramphenicol + tetracycline to select for donor and transconjugant populations, respectively. Plates were incubated overnight at 37 °C.

#### High-throughput magnetic tweezers

The magnetic tweezers apparatus is implemented on a custom-built inverted microscope described previously^63,93–95^. Briefly, collimated LED illumination (NA = 0.79, Thorlabs, Germany) passes through the gap of a neodymium magnet pair (W-05-G, SuperMagnete, Switzerland), whose height and rotation are controlled by linear motors theM-126-PD1 and C-150, with C-863 controllers (Physik Instrumente, Germany). The field of view is imaged by a CFI Plan Apochromat Lambda D 60× oil-immersion objective (Nikon, Germany) mounted on a PIFOC piezo stage (P-726, E-753 controller; Physik Instrumente, Germany). Flow-cell temperature is regulated through an objective-based heating system^96^. The temperature is regulated within approximately 0.1 °C by connecting the foil to a PID temperature controller (TC200 PID controller, Thorlabs). Images are acquired by a CMOS camera (Dalsa Falcon2, Stemmer Imaging, Germany; 4096 × 3072 pixels, 6 µm pixel size). Instrument control was implemented in a custom Python software and real-time 3D bead tracking was performed using an open-source algorithm (available at http://www.github.com/jcnossen/qtrk). The detailed parameters for the tracking are provided in **Supplementary Method**.

#### DNA constructs

The transcription template was derived from pIA1437. It comprises a T7A1 promoter followed by a 29-bp sequence that specifies incorporation of UTP only at positions 2 and 30. When RNAP initiates with ApU and transcribes in the absence of UTP, it stalls following the addition of A29. Downstream, a segment of the *E. coli rpoB* gene bears ∼350 bp from the *rfa* (*waa*) operon, representing the leader region of *rfaQ* (with the native *ops* site ∼500 bp from T7A1) and the first ∼200 bp of the *rfaQ* open reading frame. An identical, second *ops* site, was inserted by site-directed mutagenesis ∼1.1 kbp downstream of T7A1. Two construct orientations were prepared: in the assisting-force (AF) configuration the digoxigenin handle is upstream of the promoter; in the opposing-force (OF) it is downstream. Constructs were assembled by restriction–ligation^97^ (details provided in **Supplementary Method**).

#### Stalled elongation complex preparation

The RNAP holoenzyme was loaded at the promoter and stalled at the A29 position as described previously ^17,92^. Briefly, wild-type *E. coli* RNAP holoenzyme (1 nM) was incubated with either DNA construct (1:1 molar ratio) in TX buffer (50 mM HEPES pH 7.5, 100 mM KGlu, 13 mM MgAc_2_, 2 mM spermidine, 1 mM DTT, 2 mM NaN_3_, 1 mg/mL BSA) supplemented with ATP, CTP, GTP (50 µM each) and ApU dinucleotide (100 µM) at 37°C for 10 min. Heparin was added to a final concentration of 100 µg/mL for 15 min to sequester the free RNAP. Aliquots were snap-frozen in liquid nitrogen and stored at −80°C.

#### Flow cell assembly, surface preparation and tethering

Flow cells were assembled as described^98^. Two glass coverslips (#1, 24 × 60 mm, Menzel GmbH, Germany) are separated by a double-layer Parafilm channel (∼40 µL volume). The bottom coverslip is coated with 0.3% w/v nitrocellulose in amyl acetate.

Polystyrene reference beads (1.1 µm; 3 µm for 10 pN experiments) were incubated in the flow cell, followed by anti-digoxigenin antibodies (50 µg/mL, 40 min), high-salt wash (TE750 + 2 mM NaN_3_), and BSA passivation (10 mg/mL, 40 min) followed by a second high-salt wash. PBS + 2 mM NaN_3_ was used to wash away reference beads and equilibrate the flow cell after high-salt washes. The following steps were all performed in TX buffer. The stalled complexes were diluted 1:40 and incubated for 30 min in the flow cell. After washing away the excess, MyOne beads (1.05 µm; M270 2.8 µm for 10 pN) were incubated, non-specific interactions removed by applying 5 pN and washed again. Tethers showing correct force-extension behavior were selected for the assay, if the bead is not bound to multiple tethers.

#### Single-molecule transcription assay

Transcription was performed at 37°C in TX under constant applied force for 1.5 h. Reactions were restarted with NTPs (1 mM each, ATP/GTP/CTP/UTP) supplemented with proteins as indicated: RfaH (5 nM–1 µM), GreA/GreB (2 µM), NusA (200 nM), NusG (1 µM), non-biotinylated *E. coli* RNAP (nbRNAP, 10 nM).

#### Single-molecule recovery assay

To assess escape from the *ops* site pause, a two-step protocol was used: (1) a loading step, consisting in a transcription assay with 1 µM RfaH for 2–20 min; and (2) a recovery step, where the flow cell is first rinsed with 20 volumes of TX buffer and subsequently injected with TX buffer supplemented with NTPs alone, NTPs + 1 µM RfaH, NTPs + 2 µM GreA, or NTPs + 10 nM nbRNAP. Measurements continued for 1 h after injection.

#### Data processing, dwell times distributions fit and processivity

Raw reference-subtracted traces were processed with custom Python software. Traces in which RNAP restarted >10 min after NTPs addition were excluded to focus on a more homogeneous population. Tracking artefacts were removed manually. The extension was converted to base pairs using the worm-like chain model (considering 0.34 nm bp^−1^ at full extension). Gaps were filled by linear interpolation with added Gaussian noise (1 SD), and traces filtered with a Kaiser–Bessel low-pass filter at 1 Hz. Dwell times (time to add 10 consecutive nucleotides) were extracted from filtered traces, excluding those with processivity < 300 bp. The ensemble of the dwell times extracted from each data set was used to generate a dwell times distribution, fitted as described in **Supplementary Methods**. For the cases where RfaH is present, the dwell times distributions refer only to dwell times proceeding after the first *ops* site on the template (after ∼500 bp).

The processivity of each RNAP was defined as the position of the last dwell time. The resulting pool of position was used to define an empirical complementary cumulative distribution, defined as the processivity profile. The error on the processivity profile was defined as the 95% interval of the binomial standard error 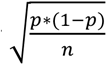 where *p* is the fraction of

RNAP that reached a certain position over the total number of RNAPs and *n* is the total number of observations.

#### Transcription time at *ops* site and analysis of the anti-pausing effect

To estimate the time spent by the RNAP at the *ops* pause site, we extracted the time needed to transcribe from 450 to 500 bp for each activity trace, given that the trace transcribed past 550 bp. From the ensemble of the times at the *ops* site, we calculated the median and the standard error on the standard deviation of the medians obtained from 1000 bootstrap procedures. The statistical significance of the difference between the distributions of the times at the *ops* site was calculated using Mann-Whitney U test and the statistical significance of the difference between the medians was defined as the 95% confidence interval bootstrapped through 100000 procedures, to calculate p-values of at least 10^-6^. Results for the statistical tests are reported in **Supplementary Tables S2-S9**.

After the *ops* site, we investigated the anti-pausing effect of NusG and RfaH, defined as a shorter median time of transcription. In detail, from the region prior the *ops* site in the condition combining RfaH, NusA and GreA, we calculated the rolling median over 10 dwell times and used the resulting median to calculate, for each trace, a 95% CI for the median from 2000 bootstrap procedures. We define an event with a shorter median transcription time the event encompassing 10 dwell times whose median is below the lower limit of the 95% CI for the same trace. For the case with NusG, the CI was extracted by bootstrapping the medians of all the region prior to the *ops* site in the condition with RfaH. We considered as a single event consecutive 10 dwell times windows that respect this condition, interrupted by at most 10 windows in the range, as a single long event can influence up to 10 windows. We considered an event to be due to RfaH if it started at most 5 dwell times after the defined *ops* site, while for NusG there was no condition on the position. The length of each event was extracted and defined as uncensored, if the end of the event happens before the end of the template, or censored, if the event starts after the *ops* site and is longer than 2 kbp. The mean and standard error of the length are calculated on the uncensored events.

#### Model for the fraction of recovery from the ops site

The recovered fraction from the ops site shows a decay with incubation time of RfaH, which was well-fitted by an exponential decay with a single rate *k*. The recovered fraction was not 1 at zero incubation time, meaning that there is a fraction of RNAPs that stalls at the *ops* site independent of RfaH *P*_stalled_. For long incubation times, the fraction of recovered RNAPs also does not decrease to zero but reaches an asymptote as a result of RNAPs that escape/bypass the *ops* site in any case *P*_bypass_. Therefore, the expression for the recovered fraction of RNAPs with incubation time is:

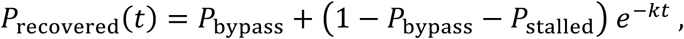

which was fit on the fraction of escaping RNAPs, with Wilson score confidence intervals to clip error values between 0 and 1, from recovery assays. The fit used the least squares method to find the local minimum to minimize the cost function (from SciPy, Python package).

## Acknowledgements

DD was supported by BaSyC – Building a Synthetic Cell” Gravitation grant (024.003.019) of the Netherlands Ministry of Education, Culture and Science (OCW) and the Netherlands Organisation for Scientific Research (NWO). IA was supported by the National Institute of General Medical Sciences (NIGMS) grant R01 GM067153. The authors used Claude Sonnet 5 (Anthropic) to assist with code development and to improve the clarity of the writing. The authors reviewed and take full responsibility for the content of this manuscript.

## Author contributions

LB, BW, IR and DD designed the project. LB performed the single-molecule experiments and analyzed the data. BW and IA provided the purified proteins. BW performed the microbiology assay. PPBA assisted with data modeling of the recovery assay. LB and DD interpreted the data. LB and DD wrote the manuscript. All authors edited the manuscript.

## Data and code availability statement

All data needed to evaluate the conclusions in the paper are present in the paper and/or the Supplementary Materials. A link to the single-molecule analysis data and scripts will be provided upon acceptance of the manuscript.

