## Supplementary Information for "RfaH licenses RNA polymerase for long-range transcription"

This document contains Supplementary Methods, 15 Supplementary figures and 12 Supplementary Tables

**Supplementary Method**

**Bead tracking algorithm**

Bead tracking is performed in real time on the CPU. For each bead, a squared region of interest (ROI) of 60 pixels is defined and a lookup table (LUT) is constructed by recording the diffraction-pattern profile over 100 axial steps of 50 nm. Axial displacement is determined by the Quadrant Interpolation (QI) algorithm using 3 iterations. Mechanical drift is corrected by an autofocus routine that repositions the piezo when the reference-bead displacement exceeds a 5 nm threshold, with a 2 s refresh interval. Data points outside the LUT range are discarded (replaced by NaN).

**PCR conditions for DNA construct preparation**

Stem fragment: amplified from pIA1437 (sequence below) by Phusion (NEB, MA, USA) using HF buffer. Cycling: 20 s initial denaturation at 98°C; 35 cycles of 10 s at 98°C, 30 s at 66°C, and extension at 72°C; 10 min final extension at 72°C.

Handle fragment: amplified from λ bacteriophage DNA by Taq (NEB, MA, USA) using Thermopol buffer in the presence of digoxigenin-conjugated UTP. Cycling: 20 s at 95°C; 35 cycles of 10 s at 95°C, 30 s at 52°C, and extension at 68°C; 10 min final extension at 68°C.

Both reactions used a Proflex PCR machine (Thermo Fisher Scientfiic, MA, USA). Primers are listed in **Supplementary Table S10**.

After BsaI-HFv2 (NEB, MA, USA) digestion and heat inactivation, fragments were purified by spin column (NEB, MA, USA) and ligated with T4 DNA ligase (NEB) in the presence of 12.5% w/v PEG-6K (Thermo Fisher Scientfiic, MA, USA). The ligation product was gel-purified from 1% agarose / 0.5× TAE stained with SYBR Safe (Thermo Fisher Scientfiic, MA, USA). Concentrations were determined by a Denovix DS-11 spectrometer (DeNovix, DE, USA) and gel images acquired on an Amersham ImageQuant 800 imaging system (Cytiva, DE, USA).

**Elongation dwell times fit-function**

The dwell times of the elongating RNAPs (**Materials and Methods**) were in the range between 0.1 s and 5400 s. The resulting distributions were fitted with a model comprising 4 functions: one gamma probability density function (pdf) (fitting the peak modelling nucleotide addition, NA), two exponential pdf's (fitting the shoulders representing short and medium pausing states, namely Pause 1 and Pause 2), and one power-law decay pdf (representing long-lived, backtrack-related pauses, LLPs). The function was fitted using simulated annealing (3000 iterations with interval update of 30) in a 100-step bootstrap procedures (constraints in **Table S11**).

$$P_{N\mathrm{nt}}(t) = \frac{f_{\mathrm{NA}}}{T_{\mathrm{NA}}(N-1)!} N(tN/T_{\mathrm{NA}})^{N-1} e^{-tN/T_{\mathrm{NA}}} + Q(t)\left[\frac{f_{\mathrm{pause\ 1}}}{T_{\mathrm{pause\ 1}}} e^{-\frac{t-T_{\mathrm{NA}}}{T_{\mathrm{pause\ 1}}}} + \frac{f_{\mathrm{pause\ 2}}}{T_{\mathrm{pause\ 2}}} e^{-t\frac{t-T_{\mathrm{NA}}}{T_{\mathrm{pause\ 2}}}} + \frac{f_{\mathrm{LLP}}\sqrt{1+T_{\mathrm{NA}}}}{2(1+t)^{3/2}}\right], \quad \text{(S1)}$$

with $Q(t, T_{\mathrm{NA}}) = \frac{(t/T_{\mathrm{NA}})^{N-1}}{1+(t/T_{\mathrm{NA}})^{N-1}}$.

The different distributions in the fit-function have weights $f_i$ and timescales $T_i$ with $i \in \{\mathrm{NA}, \mathrm{pause\ 1}, \mathrm{pause\ 2}, \mathrm{LLP}\}$. The approximations in the fit-function hold for separated timescales $T_i$ and incrementally decreasing weights $q_i$[1,2]

When these conditions are satisfied, the weights $f_i$ and characteristic timescales $T_i$ of the peak and the two shoulders can be expressed in terms of the single step probabilities $p_i$ and rates $k_i$ of each pathway

$$
\begin{aligned}
f_{\mathrm{NA}} &= (p_{NA})^N, \\
T_{\mathrm{NA}} &= N/k_{\mathrm{NA}}\,, \\
f_{\mathrm{pause\ 1}} &= \left(p_{\mathrm{NA}} + p_{\mathrm{pause\ 1}}\right)^N - (p_{\mathrm{NA}})^N\,, \\
T_{\mathrm{pause\ 1}} &= \frac{N p_{\mathrm{pause\ 1}}\left(p_{\mathrm{NA}} + p_{\mathrm{pause\ 1}}\right)^{N-1}}{f_{\mathrm{pause\ 1}}}\,\frac{1}{k_{\mathrm{pause\ 1}}}, \\
f_{\mathrm{pause\ 2}} &= \left(p_{\mathrm{NA}} + p_{\mathrm{pause\ 1}} + p_{\mathrm{pause\ 2}}\right)^N - \left(p_{\mathrm{NA}} + p_{\mathrm{pause\ 1}}\right)^N\,, \\
T_{\mathrm{pause\ 2}} &= \frac{N p_{\mathrm{pause\ 2}}\left(p_{\mathrm{NA}} + p_{\mathrm{pause\ 1}} + p_{\mathrm{pause\ 2}}\right)^{N-1}}{f_{\mathrm{pause\ 2}}}\,\frac{1}{k_{\mathrm{pause\ 2}}}.
\end{aligned}
\tag{S2}
$$

The fraction of long-lived pauses is simply obtained as $f_{\mathrm{LLP}} = 1 - f_{\mathrm{NA}} - f_{\mathrm{pause\ 1}} - f_{\mathrm{pause\ 2}}$. By inverting the relations in **Equation S2**, the single nucleotide rates $k_i$ and probabilities $p_i$ for each pathway $i \in \{\mathrm{NA, pause\ 1, pause\ 2}\}$ can be expressed in terms of the dwell-time weights $f_i$ and characteristic timescales $T_i$

$$
\begin{aligned}
p_{\mathrm{NA}} &= (f_{\mathrm{NA}})^{1/N}, \\
k_{\mathrm{NA}} &= N/T_{\mathrm{NA}}, \\
p_{\mathrm{pause\ 1}} &= \left(f_{\mathrm{NA}} + f_{\mathrm{pause\ 1}}\right)^{1/N} - (f_{\mathrm{NA}})^{1/N}, \\
k_{\mathrm{pause\ 1}} &= \frac{N p_{\mathrm{pause\ 1}}\left(p_{\mathrm{NA}} + p_{\mathrm{pause\ 1}}\right)^{N-1}}{f_{\mathrm{pause\ 1}} T_{\mathrm{pause\ 1}}}, \\
p_{\mathrm{pause\ 2}} &= \left(f_{\mathrm{NA}} + f_{\mathrm{pause\ 1}} + f_{\mathrm{pause\ 2}}\right)^{1/N} - \left(f_{\mathrm{NA}} + f_{\mathrm{pause\ 1}}\right)^{1/N}, \\
k_{\mathrm{pause\ 1}} &= \frac{N p_{\mathrm{pause\ 2}}\left(p_{\mathrm{NA}} + p_{\mathrm{pause\ 1}} + p_{\mathrm{pause\ 2}}\right)^{N-1}}{f_{\mathrm{pause\ 2}}\, T_{\mathrm{pause\ 2}}}.
\end{aligned}
\tag{S3}
$$

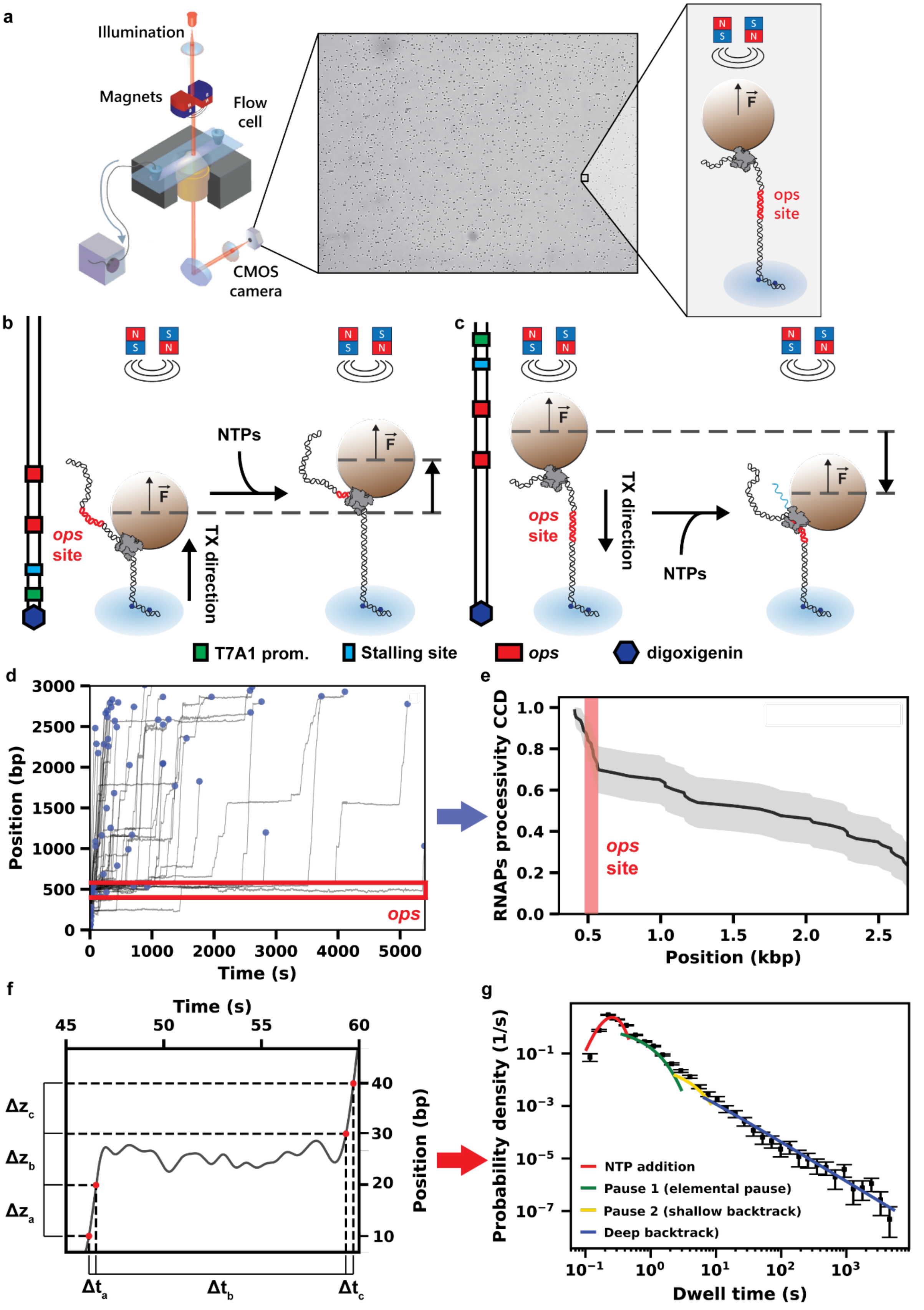


**Supplementary Figure 1: Description of the high-throughput magnetic tweezers transcription assay and data analysis.** **(a)** Simplified schematic of the magnetic tweezers instrument and typical field of view at the start of the experiment. **(b)** Schematics of the magnetic tweezers transcription assay for the case where the force is applied in the same direction as transcription elongation or **(c)** in the opposite direction. **(d)** Example of the activity traces obtained in a single experiment, after conversion of the signal (distance in nm) in base pairs (bp) and filtering to 1 Hz using a low-pass Kaiser-Bessel filter, from which the maximum positions reached (blue dots) were extracted to obtain **(e)** the complementary cumulative distribution (CCD) of the processivity of the RNAPs. The pause induced by the *ops* site is highlighted by the red flanking lines in **(d)** and semi-transparent patch at position ~500 bp in **(e)**. **(f)** Example of dwell times $\Delta t_x$, extracted as the time to add 10 consecutive nucleotides $\Delta z_x$, represented in red on a zoomed-in, low-pass filtered trace showing rapid elongation interrupted by a pause. The ensemble of the dwell times is used to generate **(g)** a dwell times distribution, with error bars deriving from 100 bootstraps procedures, on which 4 distributions are fitted: a gamma, representing NTP addition, two exponentials, describing the elemental pause and short backtracks, and a power law with exponent $t^{-3/2}$, for rare, long-lived backtracking events.

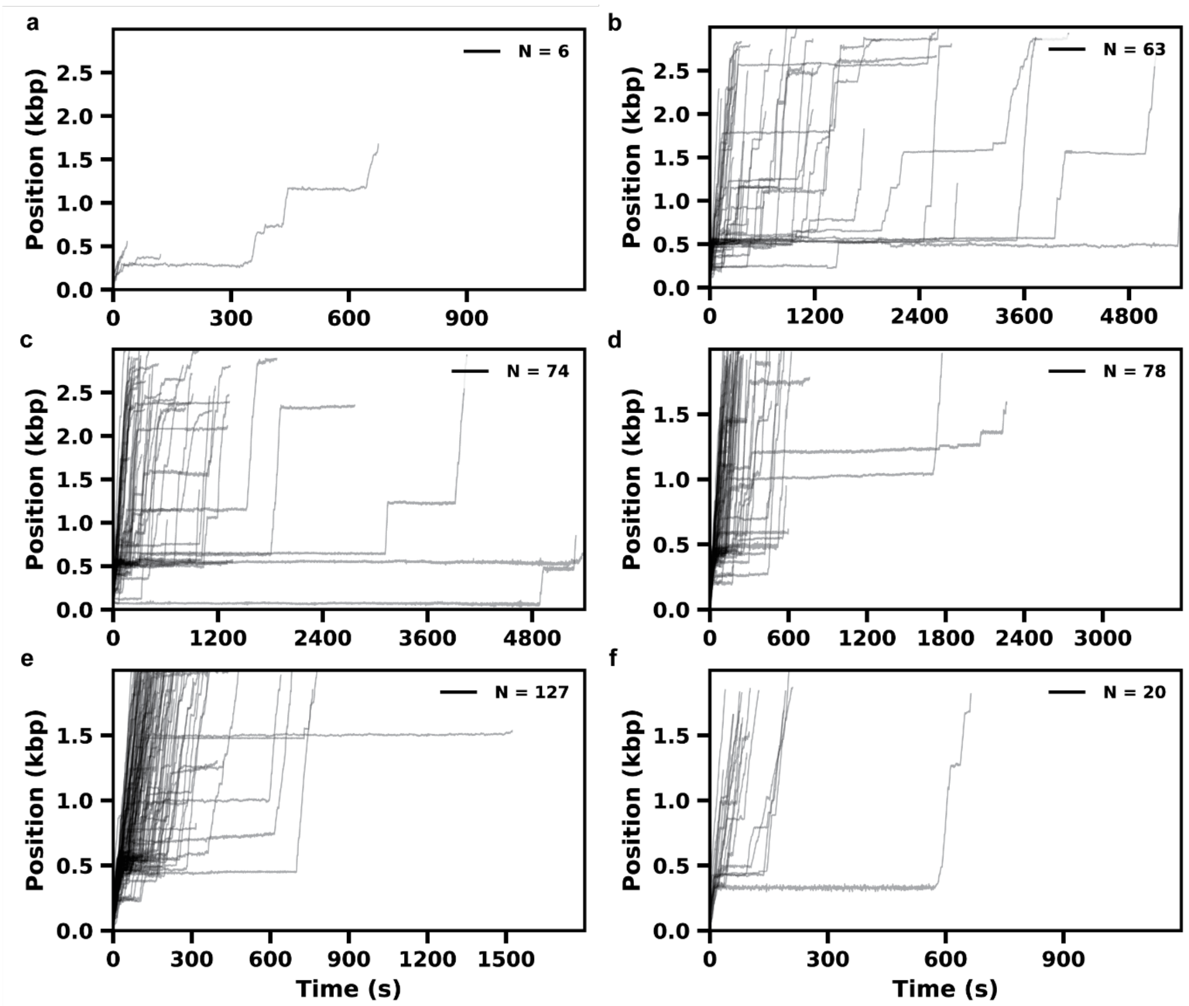


**Supplementary Figure 2: RNAP activity traces in OF and AF configurations. (a)** 20 min of activity traces under 10 pN in OF configuration. **(b)** 90 min of activity traces under 7 pN in OF configuration. **(c)** 90 min of activity traces under 3 pN in OF configuration. **(d)** 60 min of activity traces under 3 pN in AF configuration. **(e)** 30 min of activity traces under 3 pN in AF configuration. **(f)** 20 min of activity traces under 10 pN in AF configuration. All traces are filtered to 1 Hz using a low-pass Kaiser-Bessel filter.

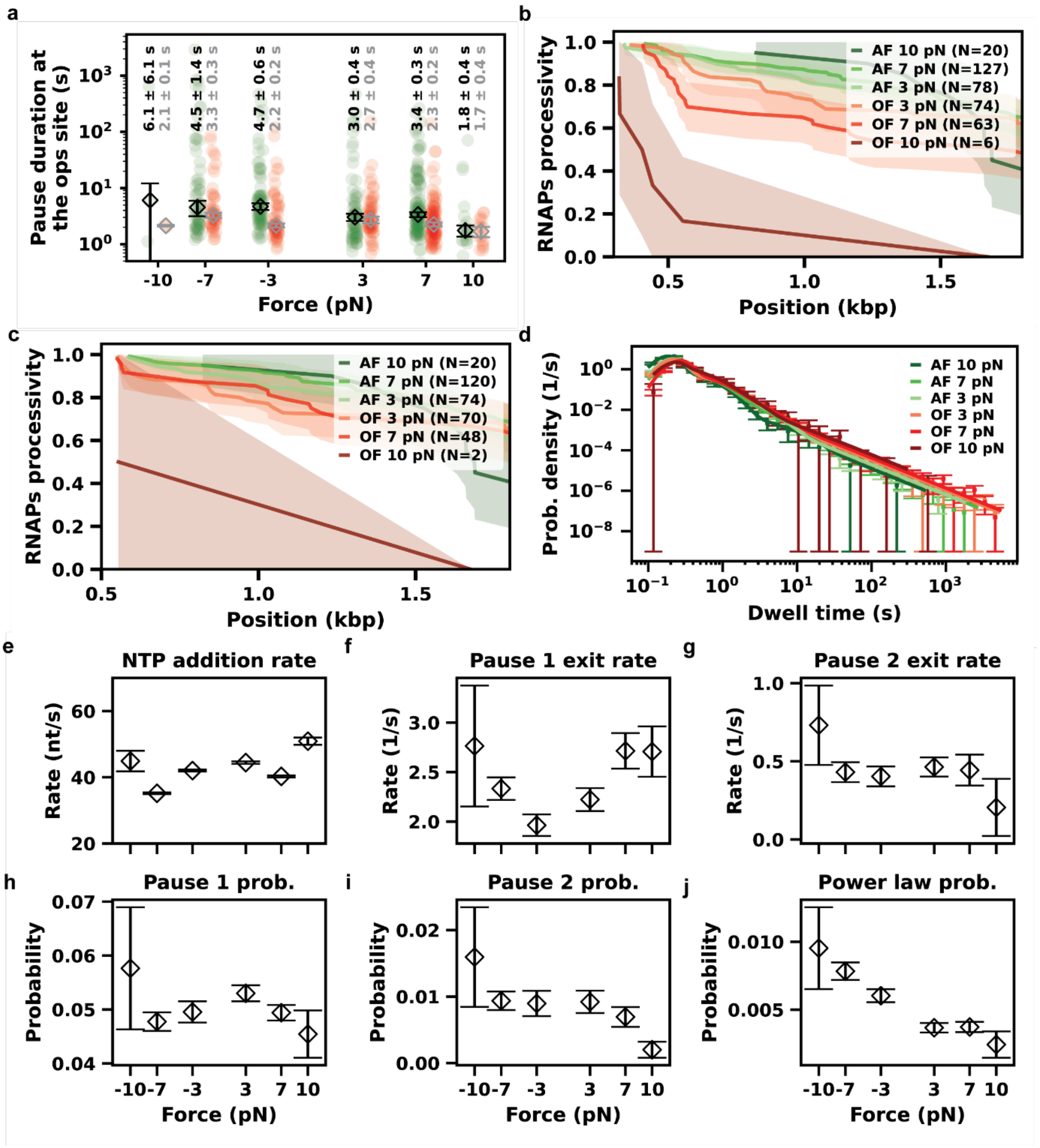


**Supplementary Figure 3: Processivity and dwell times distribution and fitting parameters for a range of forces in both AF and OF configurations. (a)** Pause durations at *ops1* (green dots) or ops2 (red dots) site for varying forces (3 to 10 pN) in both AF and OF configurations. The diamonds (black for ops1, grey for ops2) indicate the median of the respective populations, with the error bars are calculated as standard deviation of the medians from 300 bootstrap procedures (values for each population indicated above the relative scatter plot). **(b)** Processivity of the RNAPs under for varying forces (3 to 10 pN) in both AF and OF configurations for traces longer than 300 bp or **(c)** 550 bp. **(d)** Dwell times distributions for conditions in both AF and OF configurations, with error bars representing the standard deviation of the values obtained by 100 bootstrap procedures. **(e-j)** Comparison of the parameters of the dynamics of the RNAP extracted from the fit of the dwell times distribution. The error bars represent the standard deviation of the parameters obtained by the 100 bootstrap procedures.

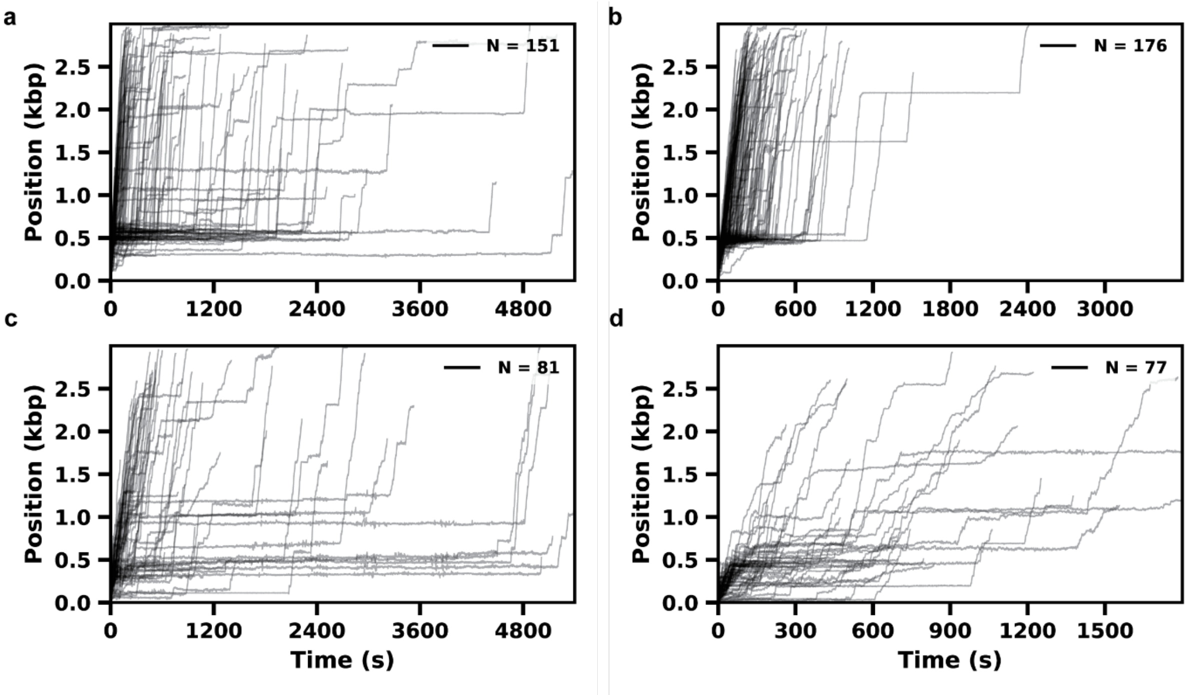


**Supplementary Figure 4: RNAP activity traces under 7 pN of applied force in OF configuration, in presence of either GreA or GreB or NusA or NusG. (a)** 90 min of activity with 2 µM GreA. **(b)** 60 min of activity with 2 µM GreB. **(c)** 90 min of activity with 1 µM NusG. All traces are filtered to 1 Hz using a low-pass Kaiser-Bessel filter. **(d)** 30 min of activity with 200 nM NusA.

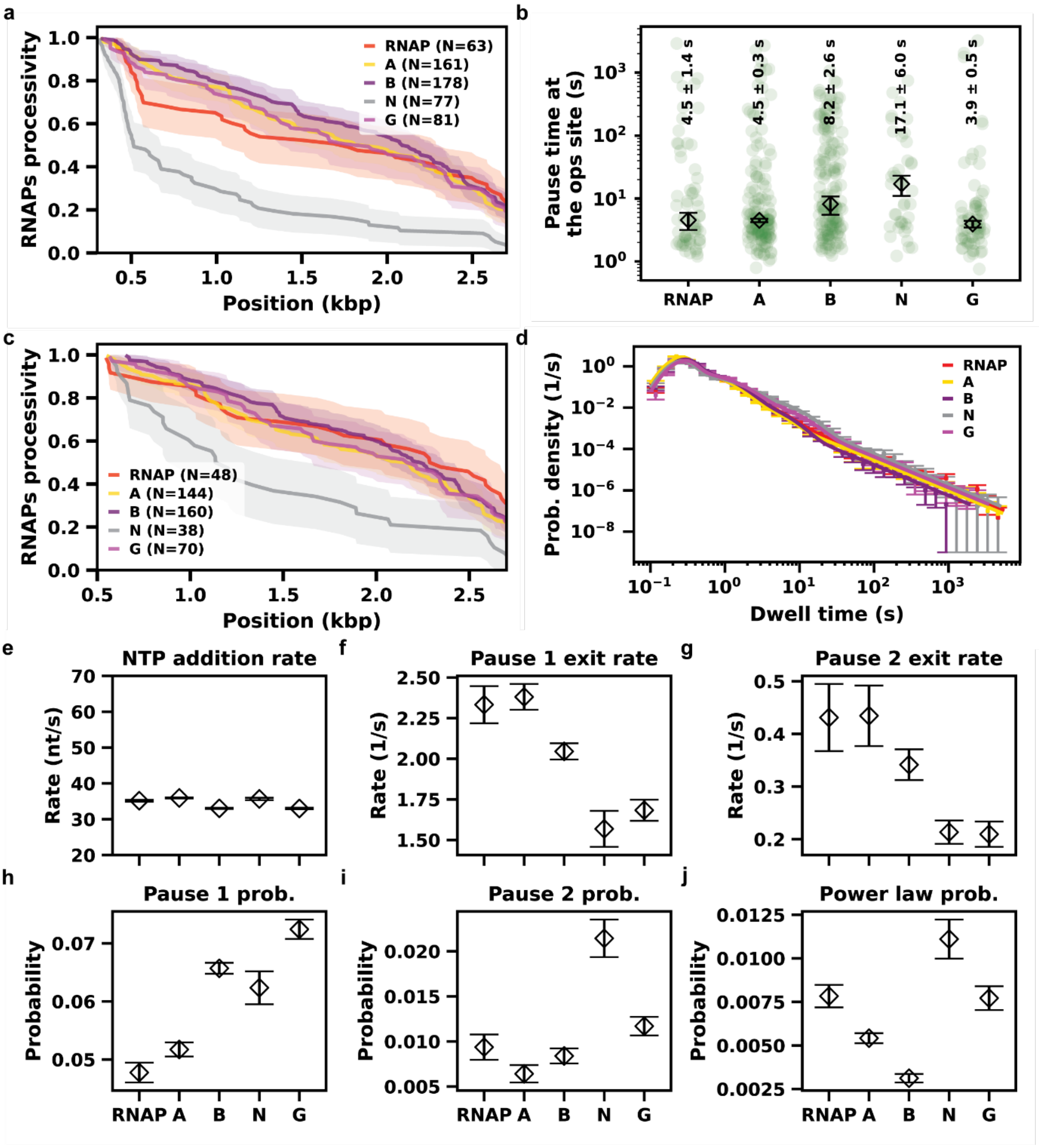


**Supplementary Figure 5: Processivity and dwell times distribution and fitting parameters under 7 pN of applied force in OF configuration, in presence of either GreA (A) or GreB (B) or NusA (N) or NusG (G). (a)** Processivity of the RNAPs under a constant force of 7 pN in OF, in the presence of the transcription factors GreA, GreB, NusG, NusA for traces longer than 300 bp. **(b)** Pause durations at *ops1* (green dots) in the absence or presence of either 2 µM GreA or 2 µM GreB or 200 nM NusA or 1 µM NusG. The diamonds indicate the median of the respective populations, with the error bars are calculated as standard deviation of the medians from 300 bootstrap procedures (values for each population indicated above the relative scatter plot). **(c)** Processivity of the RNAPs under a constant force of 7 pN in OF, in the presence of the transcription factors GreA, GreB, NusG, NusA for traces longer than 550 bp. **(d)** Dwell times distributions in the absence or presence of either 2 µM GreA or 2 µM GreB or 200 nM NusA or 1 µM NusG, with error bars representing the standard deviation of the values obtained by 100 bootstrap procedures. **(e-j)** Comparison of the parameters of the dynamics of the RNAP extracted from the fit of the dwell times distribution. The error bars represent the standard deviation of the parameters obtained by the 100 bootstrap procedures.

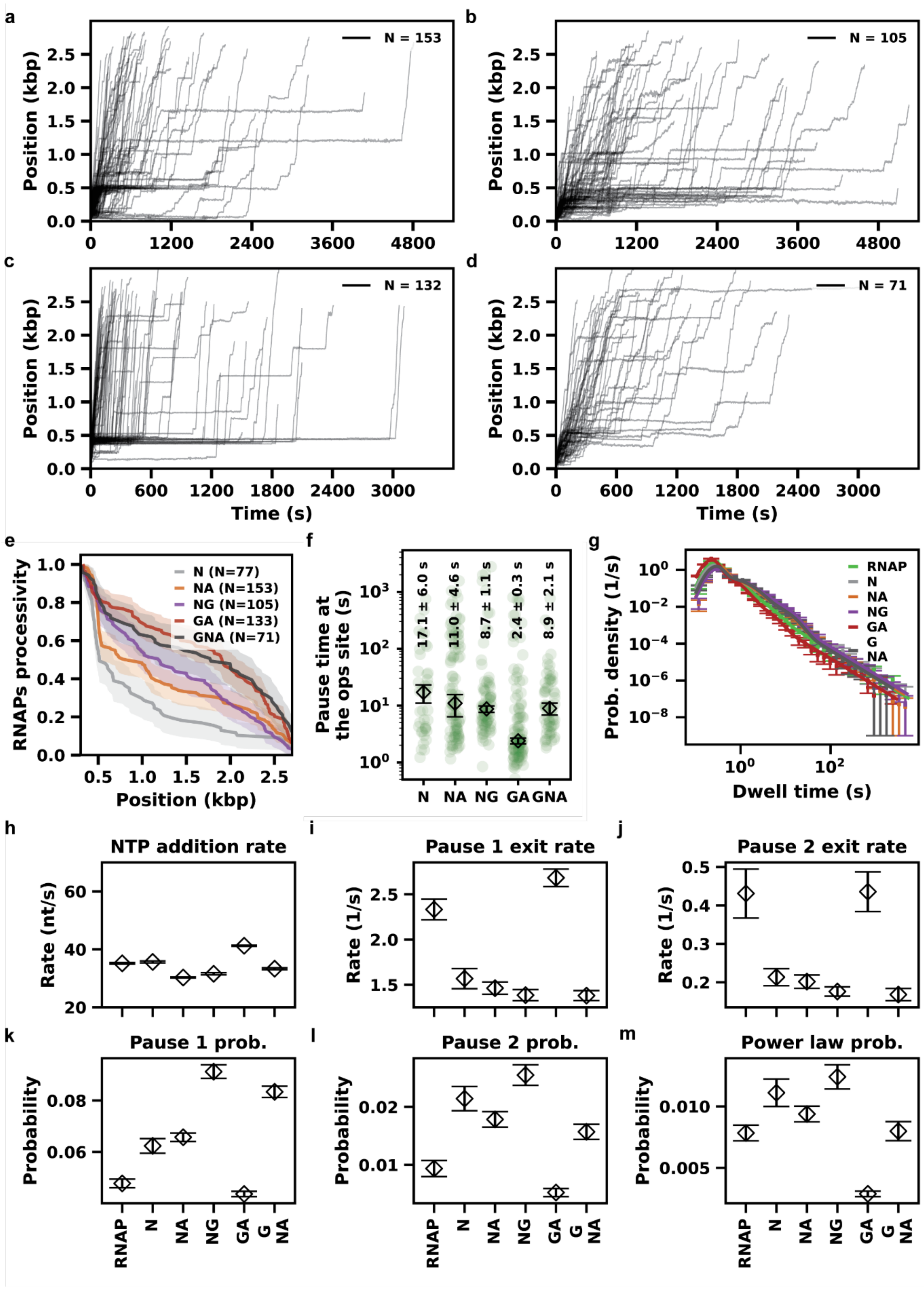


**Supplementary Figure 6: RNAP activity traces, processivity and dwell time distributions and parameters at 7 pN in OF configuration, in presence of NusA (N) and GreA (A) and/or NusG (G). (a)** 90 min of activity with 200 nM NusA + 2 µM GreA. **(b)** 90 min of activity with 200 nM NusA + 1 µM NusG. **(c)** 60 min of activity with 1 µM NusG + 2 µM GreA. **(d)** 60 min of activity with 200 nM NusA + 1 µM NusG + 2 µM GreA. All traces are filtered to 1 Hz using a low-pass Kaiser-Bessel filter. **(e)** Processivity of the RNAP for 7 pN OF with NusA

(N) in combination with either GreA (NA) or NusG (GA), the combination of the latter two (GA) or the three TFs present together (GNA). **(f)** Pause durations at *ops1* (green dots) without or with 200 nM NusA and 2 μM GreA and/or 1 μM NusG, or the combination of NusG and GreA. The diamonds indicate the median of the respective populations, with the error bars are calculated as standard deviation of the medians from 300 bootstrap procedures (values for each population indicated above the relative scatter plot). **(g)** Dwell times distributions in the absence or presence of 200 nM NusA and 2 μM GreA and/or 1 μM NusG, or the combination of NusG and GreA, compared to the wild type RNAP, with error bars representing the standard deviation of the values obtained by 100 bootstrap procedures. **(h-m)** Comparison of the parameters of the dynamics of the RNAP extracted from the fit of the dwell times distribution. The error bars represent the standard deviation of the parameters obtained by the 100 bootstrap procedures.

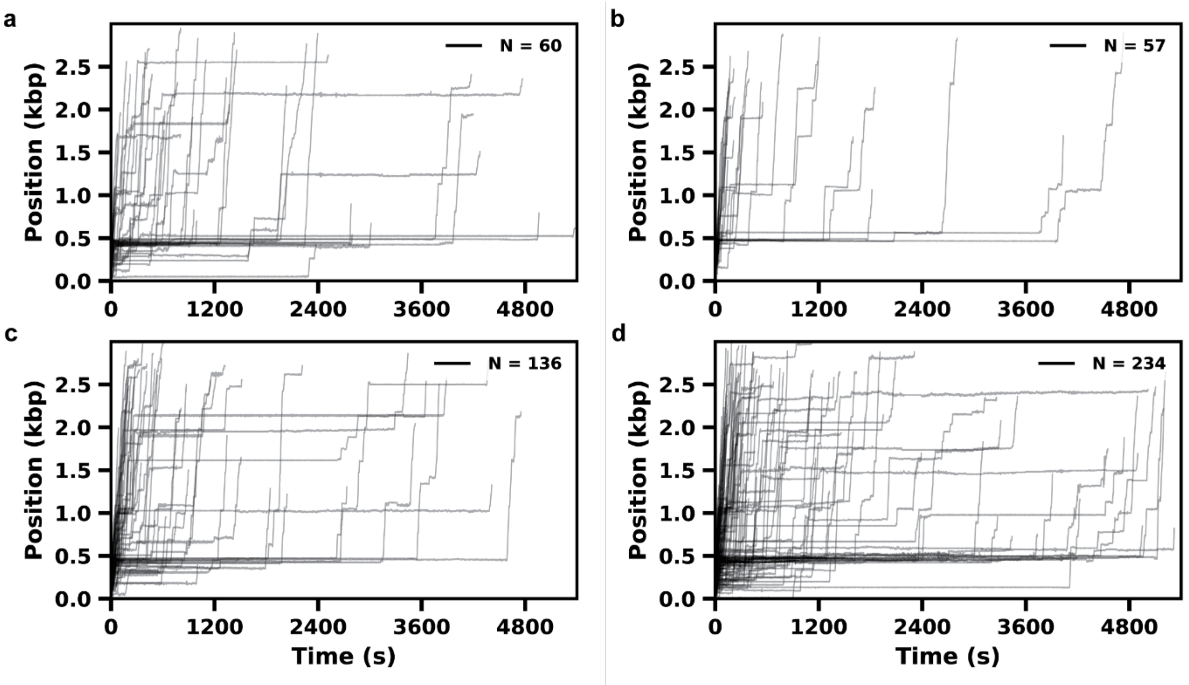


**Supplementary Figure 7: RNAP activity traces at 7 pN in OF configuration, in presence of increasing concentration of RfaH. (a)** 90 min of activity with 5 nM RfaH. **(b)** 90 min of activity with 50 nM RfaH. **(c)** 90 min of activity with 250 nM RfaH. **(d)** 90 min of activity with 1 µM RfaH. All traces are filtered to 1 Hz using a low-pass Kaiser-Bessel filter.

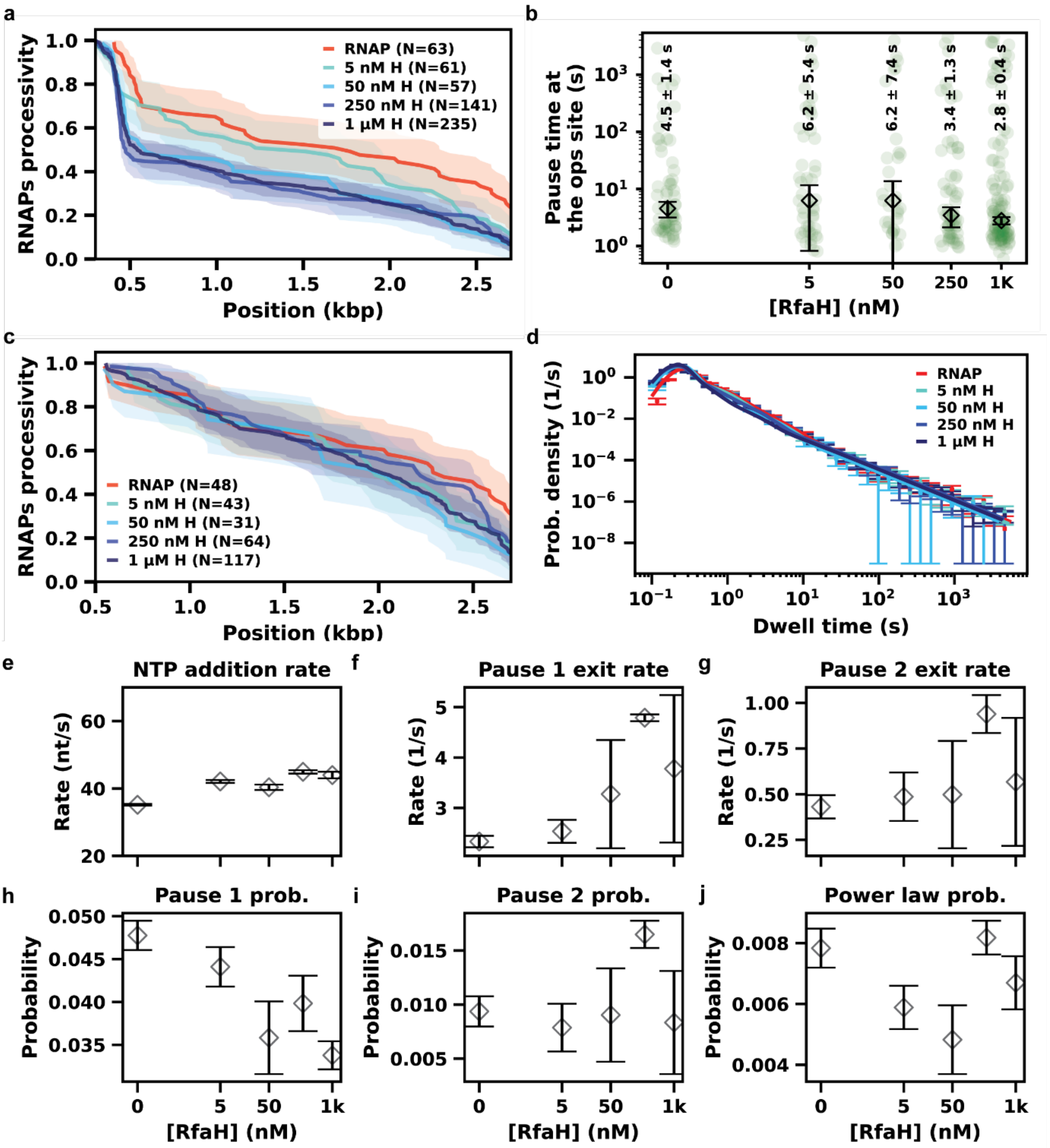


**Supplementary Figure 8: Processivity and dwell times distributions and fit parameters at 7 pN in OF configuration, in presence of increasing concentration of RfaH (H). (a)** Processivity of the RNAPs under a constant force of 7 pN in OF, in the presence of increasing concentrations of RfaH (0-1 µM) for traces longer than 300 bp. **(b)** Pause durations at *ops1* (green dots) for increasing concentrations of RfaH (0-1 µM). The diamonds indicate the median of the respective populations, with the error bars are calculated as standard deviation of the medians from 300 bootstrap procedures (values for each population indicated above the relative scatter plot). **(c)** Processivity of the RNAPs under a constant force of 7 pN in OF, in the presence of increasing concentrations of RfaH (0-1 µM) for traces longer than 550 bp. **(d)** Dwell times distributions for increasing concentrations of RfaH (0-1 µM), with error bars representing the standard deviation of the values obtained by 100 bootstrap procedures. For the conditions where RfaH is present, only dwell times after the *ops* site have been considered. **(e-j)** Comparison of the parameters of the dynamics of the RNAP extracted from the fit of the dwell times distribution. The error bars represent the standard deviation of the parameters obtained by the 100 bootstrap procedures

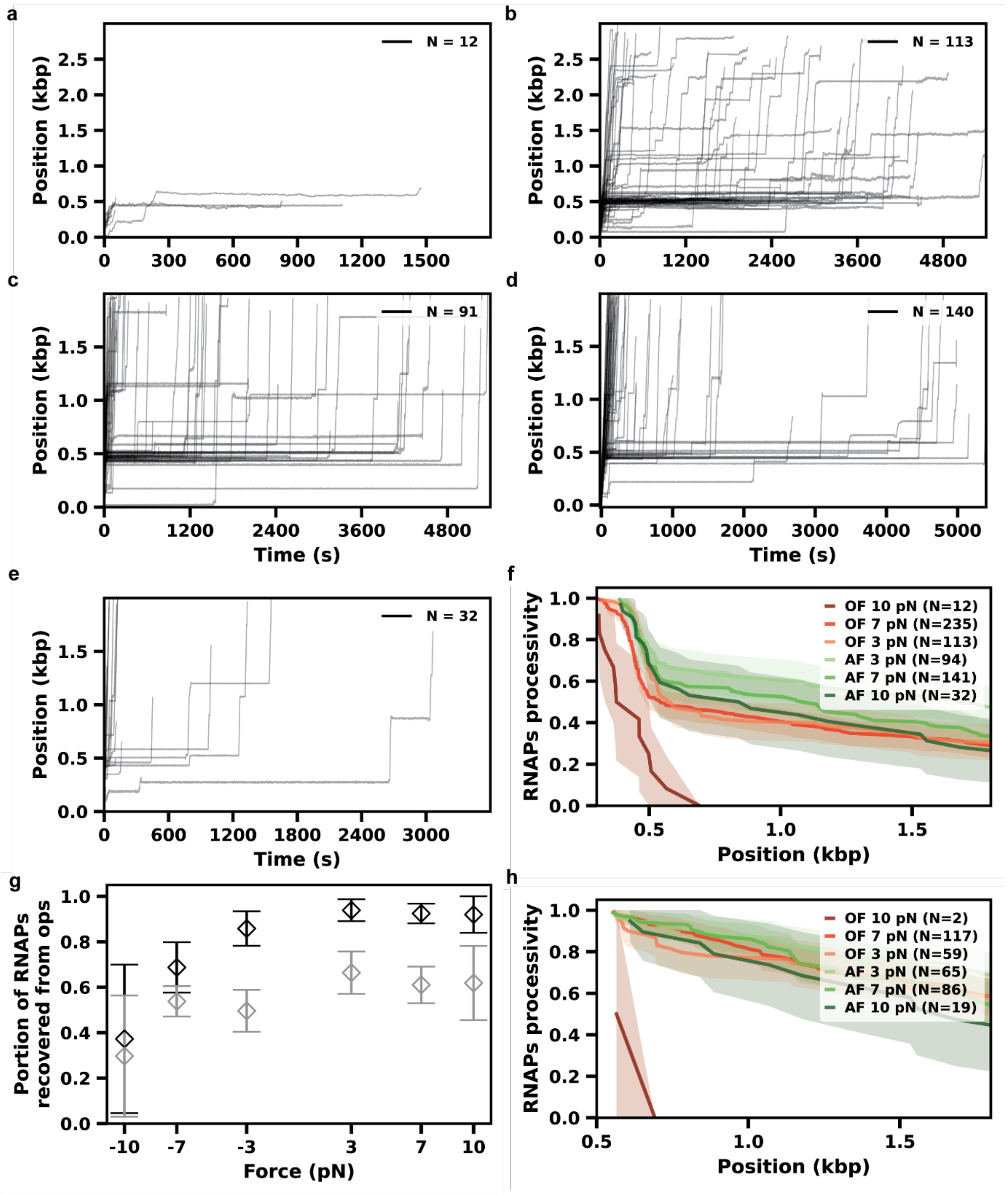


**Supplementary Figure 9: RNAP activity traces in OF and AF configuration, in presence of 1 µM RfaH, and relative processivity. (a)** 30 min of activity under 10 pN in OF configuration with 1 µM RfaH. **(b)** 90 min of activity under 3 pN in OF configuration with 1 µM RfaH. **(c)** 90 min of activity under 3 pN in AF configuration with 1 µM RfaH. **(d)** 90 min of activity under 7 pN in AF configuration with 1 µM RfaH. **(e)** 60 min of activity under 10 pN in AF configuration with 1 µM RfaH. All traces are filtered to 1 Hz using a low-pass Kaiser-Bessel filter. **(f)** Processivity of the RNAPs under for varying forces (3 to 10 pN) in both AF and OF configurations, with µM RfaH, for traces longer than 300 bp. **(g)** Comparison of the fraction of RNAPs that recover from the *ops* site for varying forces (3 to 10 pN) in both OF and AF configurations without (black) or with 1 µM RfaH (grey). The errors bars represent the Wilson score with 95% CI. **(h)** Processivity of the RNAPs under for varying forces (3 to 10 pN) in both AF and OF configurations, with µM RfaH, for traces longer than 300 bp.

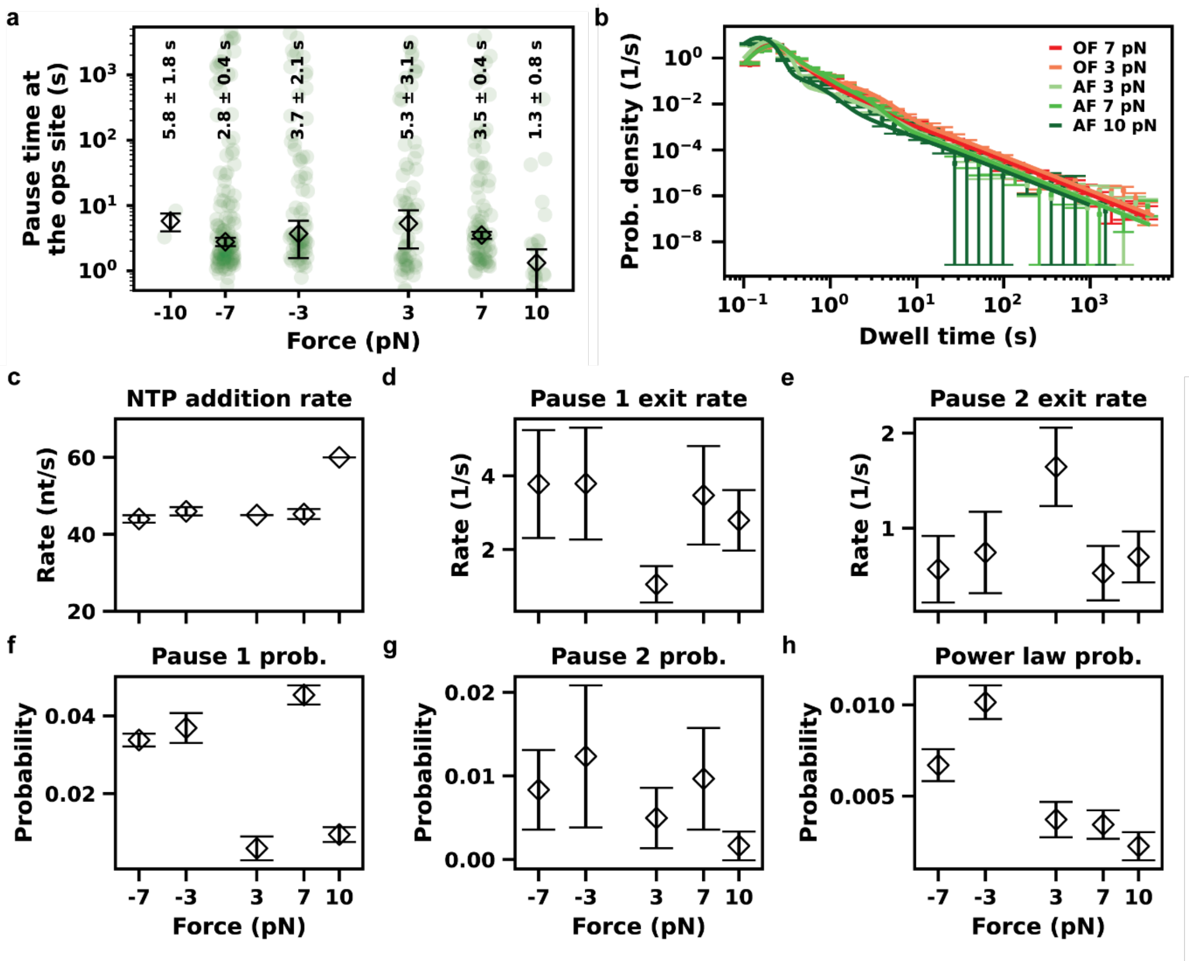


**Supplementary Figure 10. Dwell times distribution and fitting parameters for a range of forces in both AF and OF configurations, in the presence of RfaH. (a) )** Pause durations at *ops1* (green dots) for varying force (3 to 10 pN) in both OF and AF configurations in the presence of 1 µM RfaH. The diamonds indicate the median of the respective populations, with the error bars are calculated as standard deviation of the medians from 300 bootstrap procedures (values for each population indicated above the relative scatter plot). **(b)** Dwell times distributions for varying forces (3 to 10 pN) in both AF and OF configurations after the *ops* site in presence of 1 µM RfaH, with error bars representing the standard deviation of the values obtained by 100 bootstrap procedures. **(c-h)** Comparison of the parameters of the dynamics of the RNAP extracted from the fit of the dwell times distribution. The error bars represent the standard deviation of the parameters obtained by the 100 bootstrap procedures.

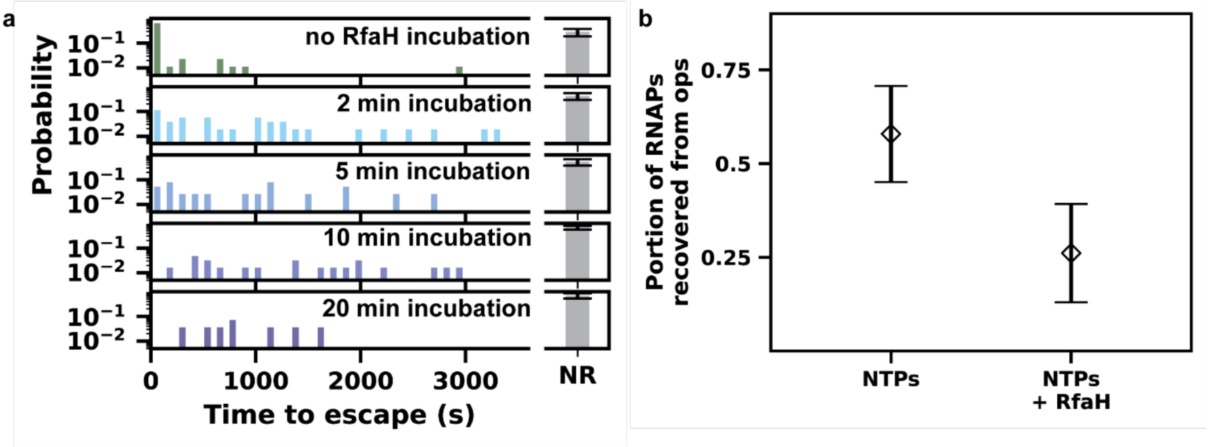


**Supplementary Figure 11. (a)** Histogram (2 min bins) of time to escape from *ops* during recovery assay after no incubation with RfaH or 2, 5, 10 or 20 min of incubation with 1 µM RfaH, resuming transcription with NTPs only. The grey bars represent the stalled population, and the error bars are the Wilson score with 95% CI. **(b)** Fraction of RNAPs that escape from the *ops* site in the recovery assay after 10 min incubation of RfaH, resuming transcription with either NTPs only or with 1 µM RfaH. The error bars represent the Wilson score with 95% CI.

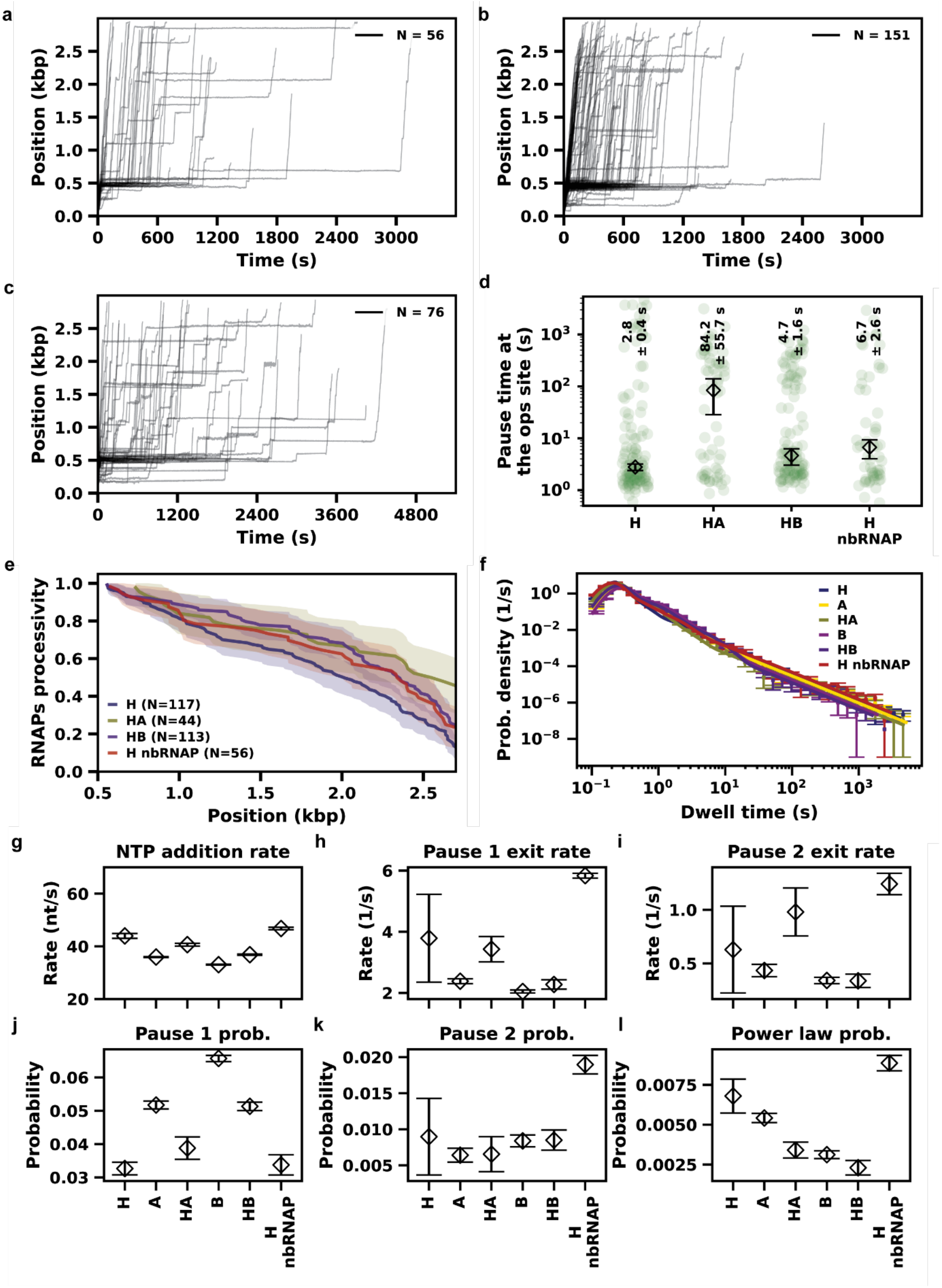


**Supplementary Figure 12: RNAP activity traces under 7 pN of applied force in OF configuration, in presence of RfaH (H) and either GreA (A) or GreB (B) or non-biotinylated RNAP (nbRNAP), with related dwell times distributions and fit parameters. (a)** 60 min of activity with 1 µM RfaH + 2 µM GreA. **(b)** 60 min of activity with 1 µM RfaH + 2 µM GreB. **(c)** 90 min of activity with 1 µM RfaH + 10 nM nbRNAP. All traces are filtered to 1 Hz using a low-pass Kaiser-Bessel filter. **(d)** Pause durations at *ops1* (green dots) in the presence of 1 μM RfaH alone or with either 2 μM GreA or 2 μM GreB or 10 nM nbRNAP. The diamonds indicate the median of the respective populations, with the error bars are calculated as standard deviation of the medians from 300 bootstrap procedures (values for each population indicated above the relative scatter plot). **(e)** Processivity of the RNAPs under 7 pN of applied force in OF configuration, with 1 μM RfaH in isolation or in combination with either 2 μM GreA or 2 μM GreB or 10 nM nbRNAP, for traces longer than 550 bp. **(f)** Dwell times distributions in the presence of 1 μM RfaH alone or with either 2 μM GreA or 2 μM GreB or 10 nM nbRNAP, compared with the individual TFs in isolation, with error bars representing the standard deviation of the values obtained by 100 bootstrap procedures. For the conditions where RfaH is present, only dwell times after the *ops* site have been considered. **(g-l)** Comparison of the parameters of the dynamics of the RNAP extracted from the fit of the dwell times distribution. The error bars represent the standard deviation of the parameters obtained by the 100 bootstrap procedures.

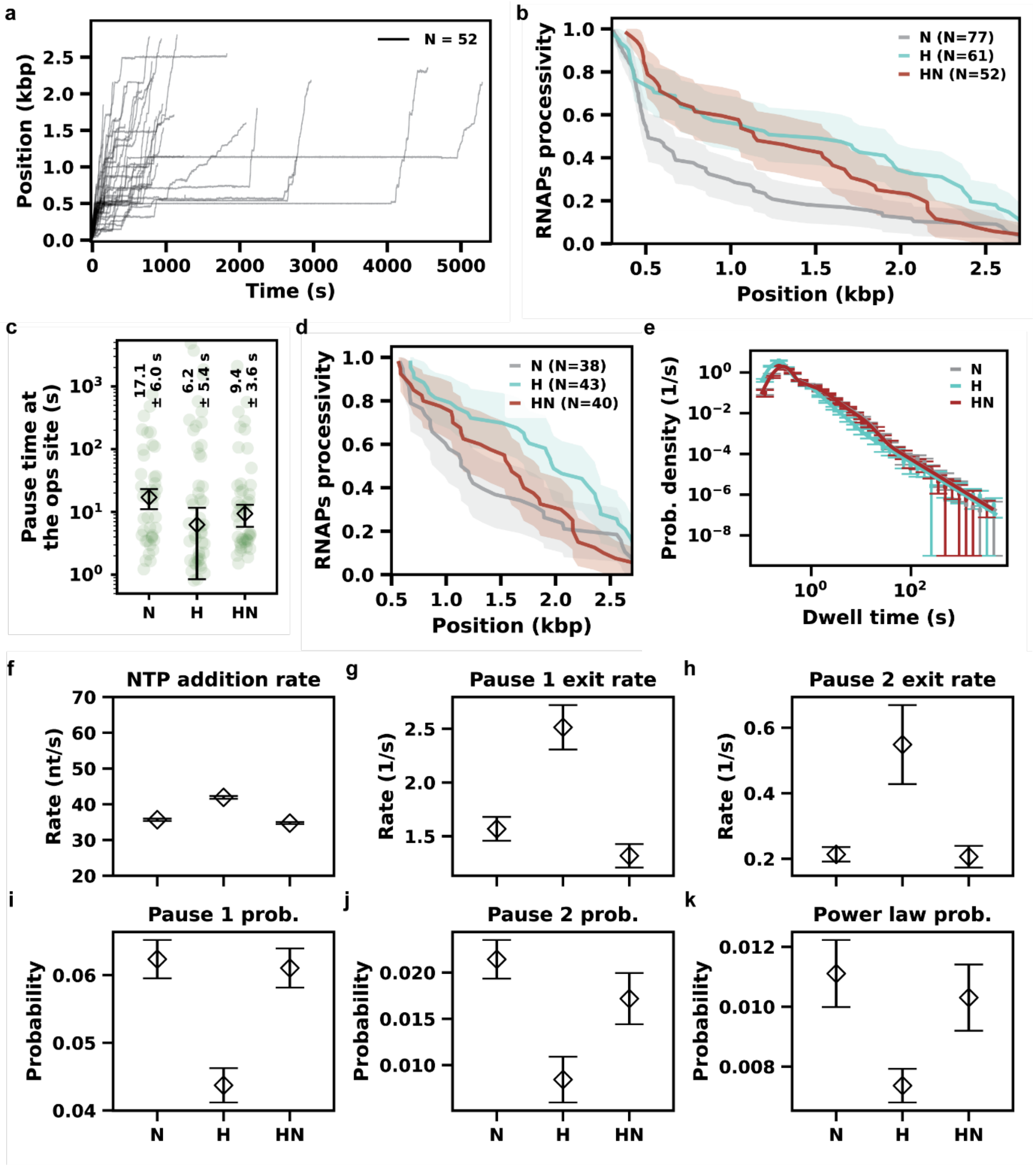


**Supplementary Figure 13: RNAP activity traces, processivity and dwell time distributions and parameters under 7 pN of applied force in OF configuration, in presence of NusA (N) and RfaH (H). (a)** First 90 min of activity with 5 nM RfaH + 200 nM NusA**.** All traces are filtered to 1 Hz using a low-pass Kaiser-Bessel filter. **(b)** Processivity of the RNAPs under 7 pN in OF configuration, in the presence of 200 nM NusA and/or 5 nM RfaH, for traces longer than 300 bp. **(c)** Pause durations at *ops1* (green dots) with 200 nM NusA and/or 5 nM RfaH. The diamonds indicate the median of the respective populations, with the error bars are calculated as standard deviation of the medians from 300 bootstrap procedures (values for each population indicated above the relative scatter plot). **(d)** Processivity of the RNAPs under 7 pN in OF configuration, in the presence of 200 nM NusA and/or 5 nM RfaH, for traces longer than 550 bp. **(e)** Dwell times distributions in the absence or presence of 200 nM NusA and/or 5 nM RfaH, with error bars representing the standard deviation of the values obtained by 100 bootstrap procedures. For the conditions where RfaH is present, only dwell times after the *ops* site have been considered. **(f-k)** Comparison of the parameters of the dynamics of the RNAP extracted from the fit of the dwell times distribution. The error bars represent the standard deviation of the parameters obtained by the 100 bootstrap procedures.

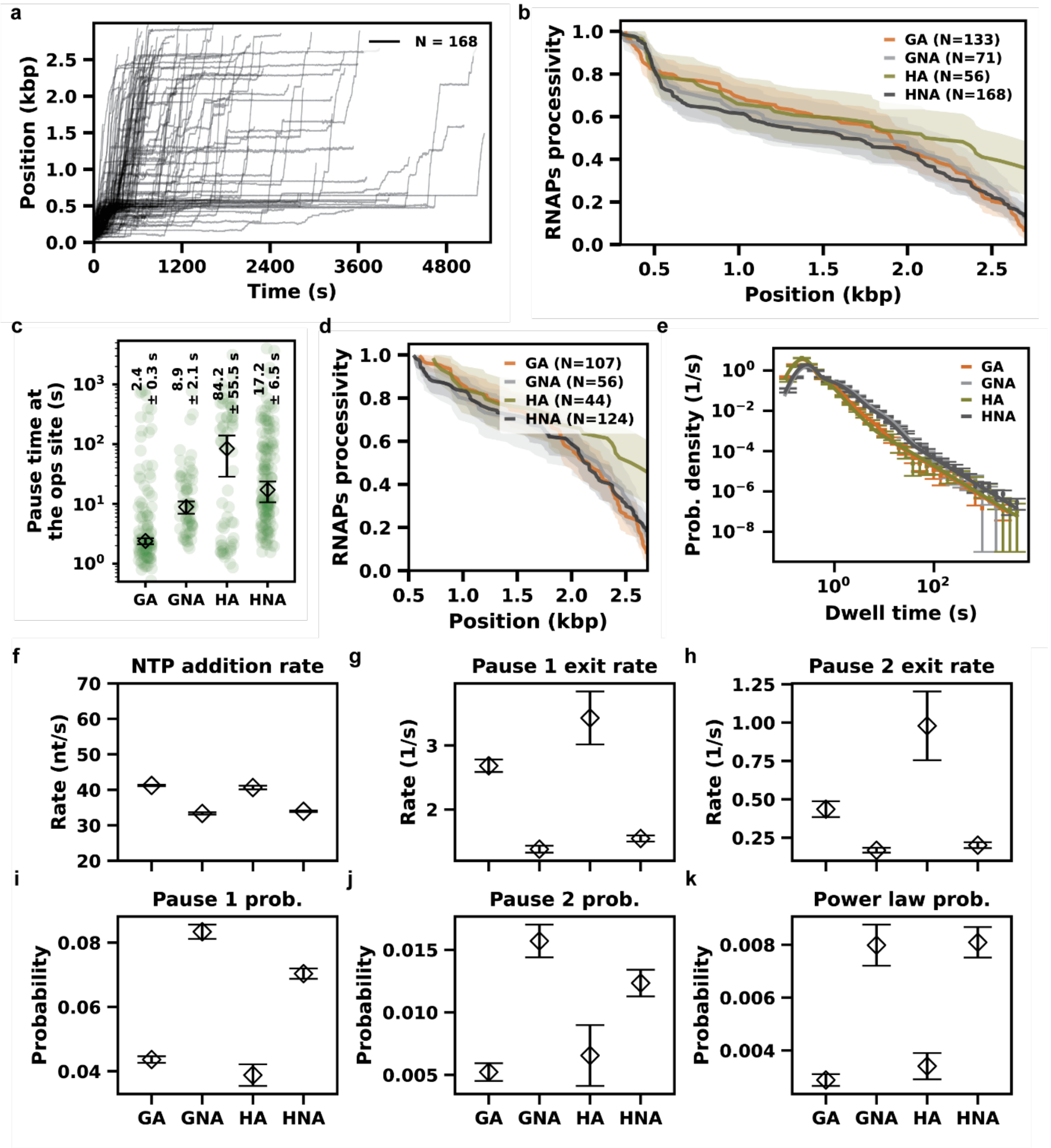


**Supplementary Figure 14: RNAP activity traces, processivity and dwell time distributions and parameters in presence of GreA (A), NusA (N) and RfaH (H), compared to NusG (G). (a)** 90 min of activity with 1 µM RfaH + 2 µM GreA + 200 nM NusA. All traces are filtered to 1 Hz using a low-pass Kaiser-Bessel filter. **(b)** Processivity of the RNAPs under 7 pN in OF configuration, in the presence of 2 µM GreA and either 1 µM RfaH or 1 µM NusG, without or with 200 nM NusA, for traces longer than 300 bp. **(c)** Pause durations at *ops1* (green dots) in the presence of 2 µM GreA and either 1 µM RfaH or 1 µM NusG, without or with 200 nM NusA. The diamonds indicate the median of the respective populations, with the error bars are calculated as standard deviation of the medians from 300 bootstrap procedures (values for each population indicated above the relative scatter plot). **(d)** Processivity of the RNAPs under 7 pN in OF configuration, in the presence of 2 µM GreA and either 1 µM RfaH or 1 µM NusG, without or with 200 nM NusA, for traces longer than 550 bp. **(e)** Dwell times distributions in the presence of 2 µM GreA and either 1 µM RfaH or 1 µM NusG, without or with 200 nM NusA, with error bars representing the standard deviation of the values obtained by 100 bootstrap procedures. **(f-k)** Comparison of the parameters of the dynamics of the RNAP extracted from the fit of the dwell times distribution. The error bars represent the standard deviation of the parameters obtained by the 100 bootstrap procedures.

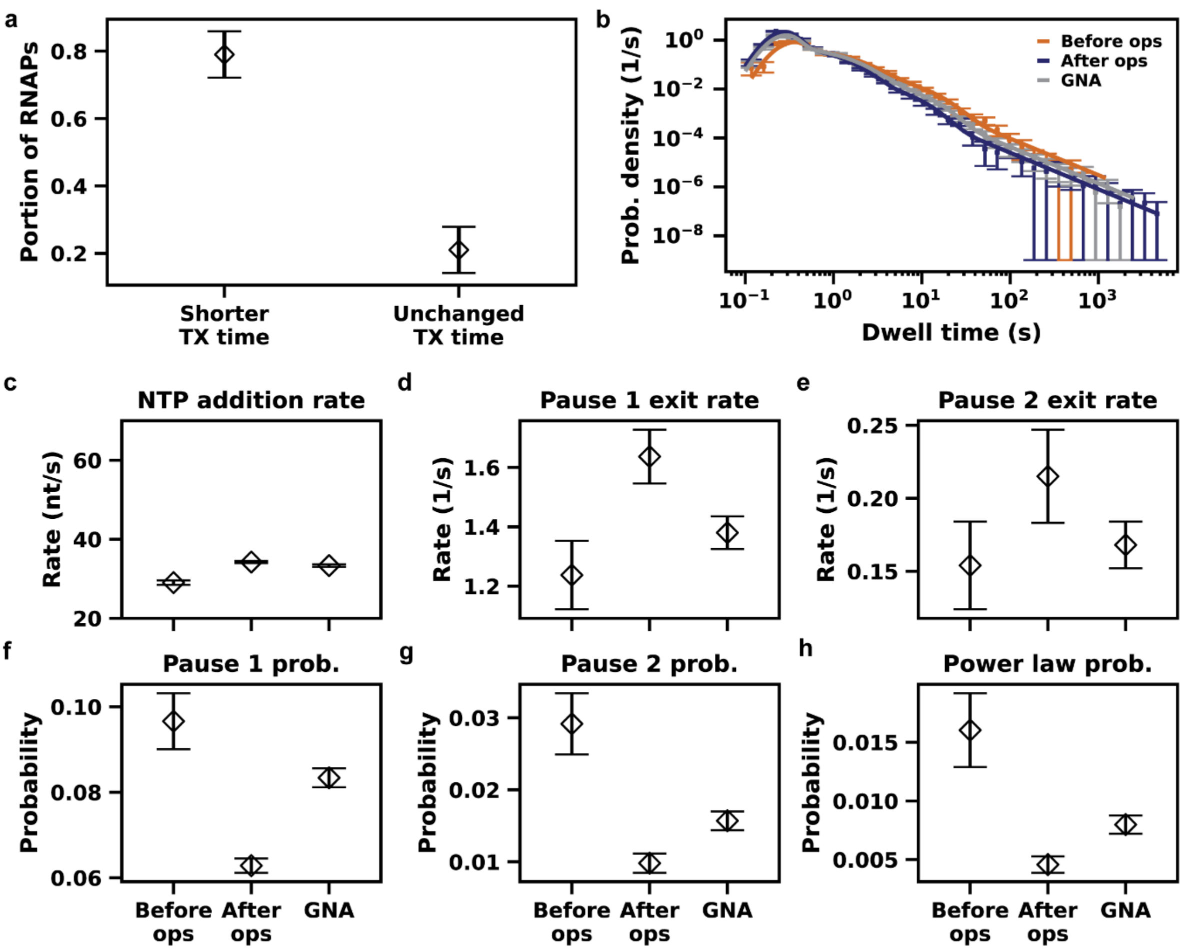


**Supplementary Figure 15: Dwell time distributions and parameters in presence of RfaH, GreA and NusA, comparing the dynamics before the *ops* site or after it, for regions with lower median transcription time.** **(a)** Portion of the RNAPs showing a median transcription (TX) time lower or above the lower end of the 95% CI of the medians over a window of 10 dwell times obtained from 2000 bootstrap procedures.. **(b)** Comparison of the dwell times distributions in the presence of 1 µM RfaH, 2 µM GreA and 200 nM NusA, before the *ops* site or after it, in the latter case for the regions with median TX time below the lower end of the 95% CI of the medians over a window of 10 dwell times, obtained from 2000 bootstrap procedures. The error bars represent the standard deviation of the values obtained by 100 bootstrap procedures. **(c-h)** Comparison of the parameters of the dynamics of the RNAP extracted from the fit of the dwell times distributions. The error bars represent the standard deviation of the parameters obtained by the 100 bootstrap procedures.

**Supplementary Table 1.** Overview of the experimental conditions explored in the present study, reporting force configuration (assisting, AF, or opposing, OF), magnitude of the force (in pN), protein concentrations (in nM), number of traces and dwell times extracted, kinetic parameters fitted (mean and standard deviation).

| ID | Force configuration | Force (pN) | [RfaH] (nM) | [NusA] (nM) | [GreA] (nM) | [GreB] (nM) | [NusG] (nM) | [nbRNAP] (nM) | N. traces | N. dwell times |
|---|---|---|---|---|---|---|---|---|---|---|
| 1 | AF | 10 | - | - | - | - | - | - | 20 | 2998 |
| 2 | AF | 7 | - | - | - | - | - | - | 106 | 16715 |
| 3 | AF | 3 | - | - | - | - | - | - | 73 | 11189 |
| 4 | OF | 3 | - | - | - | - | - | - | 74 | 13627 |
| 5 | OF | 7 | - | - | - | - | - | - | 63 | 9759 |
| 6 | OF | 10 | - | - | - | - | - | - | 6 | 404 |
| 7 | OF | 7 | - | - | 2000 | - | - | - | 161 | 26318 |
| 8 | OF | 7 | - | - | - | 2000 | - | - | 178 | 31402 |
| 9 | OF | 7 | - | 200 | - | - | - | - | 77 | 6768 |
| 10 | OF | 7 | - | - | - | - | 1000 | - | 81 | 13757 |
| 11 | OF | 7 | - | 200 | 2000 | - | - | - | 153 | 17662 |
| 12 | OF | 7 | - | 200 | - | - | 1000 | - | 105 | 13969 |
| 13 | OF | 7 | - | - | 2000 | - | 1000 | - | 133 | 19587 |
| 14 | OF | 7 | - | 200 | 2000 | - | 1000 | - | 71 | 11234 |
| 15 | OF | 7 | 5 | - | - | - | - | - | 61 | 8523 |
| 16 | OF | 7 | 50 | - | - | - | - | - | 57 | 6462 |
| 17 | OF | 7 | 250 | - | - | - | - | - | 141 | 14667 |
| 18 | OF | 7s | 1000 | - | - | - | - | - | 235 | 25586 |
| 19 | OF | 10 | 1000 | - | - | - | - | - | 12 | 503 |
| 20 | OF | 3 | 1000 | - | - | - | - | - | 113 | 13045 |
| 21 | AF | 3 | 1000 | - | - | - | - | - | 89 | 10602 |
| 22 | AF | 7 | 1000 | - | - | - | - | - | 132 | 14447 |
| 23 | AF | 10 | 1000 | - | - | - | - | - | 29 | 2451 |
| 24 | OF | 7 | 5 | 200 | - | - | - | - | 52 | 6537 |
| 25 | OF | 7 | 1000 | - | 2000 | - | - | - | 56 | 9972 |
| 26 | OF | 7 | 1000 | - | - | 2000 | - | - | 151 | 24322 |
| 27 | OF | 7 | 1000 | - | - | - | - | 10 | 76 | 11296 |
| 28 | OF | 7 | 1000 | 200 | 2000 | - | - | - | 168 | 25175 |

| ID | NA rate (nt/s, mean) | NA rate (nt/s, std) | P1 rate (1/s, mean) | P1 rate (1/s, std) | P2 rate (1/s, mean) | P2 rate (1/s, std) | P1 prob. (mean) | P1 prob. (std) | P2 prob. (mean ) | P2 prob. (std) | BT prob. (mean) | BT prob. (std) |
|---|---|---|---|---|---|---|---|---|---|---|---|---|
| 1 | 50.9054 | 1.1204 | 2.7069 | 0.2546 | 0.2054 | 0.1829 | 0.0454 | 0.0044 | 0.0020 | 0.0012 | 0.0024 | 0.0010 |
| 2 | 40.2704 | 0.2911 | 2.7137 | 0.1801 | 0.4435 | 0.0988 | 0.0494 | 0.0014 | 0.0069 | 0.0015 | 0.0037 | 0.0004 |
| 3 | 44.4375 | 0.3728 | 2.2227 | 0.1162 | 0.4640 | 0.0614 | 0.0530 | 0.0015 | 0.0092 | 0.0017 | 0.0037 | 0.0003 |
| 4 | 42.0171 | 0.2725 | 1.9631 | 0.1087 | 0.4037 | 0.0638 | 0.0495 | 0.0020 | 0.0090 | 0.0019 | 0.0060 | 0.0005 |
| 5 | 35.1658 | 0.2554 | 2.3329 | 0.1142 | 0.4311 | 0.0637 | 0.0478 | 0.0017 | 0.0094 | 0.0014 | 0.0078 | 0.0006 |
| 6 | 44.9010 | 3.0847 | 2.7634 | 0.6113 | 0.7312 | 0.2541 | 0.0576 | 0.0113 | 0.0159 | 0.0075 | 0.0095 | 0.0030 |
| 7 | 35.9401 | 0.1548 | 2.3806 | 0.0793 | 0.4345 | 0.0575 | 0.0517 | 0.0012 | 0.0064 | 0.0010 | 0.0054 | 0.0003 |
| 8 | 33.0391 | 0.1597 | 2.0452 | 0.0499 | 0.3415 | 0.0293 | 0.0657 | 0.0009 | 0.0084 | 0.0008 | 0.0031 | 0.0002 |

| | | | | | | | | | | | |
|---|---|---|---|---|---|---|---|---|---|---|---|
| 9 | 35.6579 | 0.3458 | 1.5684 | 0.1110 | 0.2134 | 0.0222 | 0.0624 | 0.0028 | 0.0214 | 0.0021 | 0.0111 | 0.0011 |
| 10 | 32.9968 | 0.2172 | 1.6833 | 0.0653 | 0.2095 | 0.0241 | 0.0724 | 0.0017 | 0.0117 | 0.0010 | 0.0077 | 0.0007 |
| 11 | 30.2701 | 0.2155 | 1.4628 | 0.0677 | 0.2018 | 0.0175 | 0.0657 | 0.0016 | 0.0178 | 0.0013 | 0.0094 | 0.0006 |
| 12 | 31.5669 | 0.3553 | 1.3854 | 0.0615 | 0.1760 | 0.0122 | 0.0912 | 0.0026 | 0.0255 | 0.0018 | 0.0124 | 0.0010 |
| 13 | 41.2360 | 0.1991 | 2.6826 | 0.0974 | 0.4358 | 0.0517 | 0.0436 | 0.0010 | 0.0052 | 0.0007 | 0.0029 | 0.0002 |
| 14 | 33.3283 | 0.3093 | 1.3801 | 0.0552 | 0.1681 | 0.0160 | 0.0834 | 0.0022 | 0.0157 | 0.0013 | 0.0080 | 0.0008 |
| 15 | 42.0866 | 0.4302 | 2.5360 | 0.2269 | 0.4861 | 0.1328 | 0.0441 | 0.0023 | 0.0079 | 0.0022 | 0.0059 | 0.0007 |
| 16 | 40.3655 | 0.8005 | 3.2743 | 1.0747 | 0.4979 | 0.2939 | 0.0358 | 0.0042 | 0.0090 | 0.0043 | 0.0048 | 0.0011 |
| 17 | 44.9392 | 0.4867 | 4.7909 | 0.0678 | 0.9391 | 0.1039 | 0.0398 | 0.0032 | 0.0165 | 0.0013 | 0.0082 | 0.0006 |
| 18 | 43.9472 | 0.9093 | 3.7868 | 1.4371 | 0.6286 | 0.4041 | 0.0327 | 0.0019 | 0.0090 | 0.0053 | 0.0068 | 0.0011 |
| 19 | 38.3466 | 2.0267 | 1.9340 | 0.5379 | 0.3524 | 0.0862 | 0.0692 | 0.0105 | 0.0211 | 0.0067 | 0.0104 | 0.0029 |
| 20 | 45.9655 | 1.0790 | 3.7895 | 1.5185 | 0.7441 | 0.4280 | 0.0369 | 0.0038 | 0.0123 | 0.0085 | 0.0101 | 0.0009 |
| 21 | 44.9976 | 0.0036 | 1.0570 | 0.4923 | 1.6456 | 0.4128 | 0.0059 | 0.0031 | 0.0050 | 0.0036 | 0.0037 | 0.0010 |
| 22 | 45.2475 | 1.3096 | 3.4714 | 1.3364 | 0.5261 | 0.2865 | 0.0454 | 0.0025 | 0.0097 | 0.0061 | 0.0034 | 0.0008 |
| 23 | 59.9966 | 0.0105 | 2.7932 | 0.8177 | 0.6971 | 0.2681 | 0.0095 | 0.0019 | 0.0016 | 0.0017 | 0.0023 | 0.0008 |
| 24 | 34.4768 | 0.4298 | 1.4112 | 0.1216 | 0.2199 | 0.0374 | 0.0606 | 0.0035 | 0.0186 | 0.0029 | 0.0104 | 0.0013 |
| 25 | 40.6365 | 0.5168 | 3.4290 | 0.4141 | 0.9793 | 0.2240 | 0.0388 | 0.0034 | 0.0065 | 0.0024 | 0.0034 | 0.0005 |
| 26 | 36.8335 | 0.2463 | 2.2750 | 0.1522 | 0.3383 | 0.0630 | 0.0513 | 0.0012 | 0.0085 | 0.0014 | 0.0023 | 0.0005 |
| 27 | 46.7570 | 0.4626 | 5.8314 | 0.0784 | 1.2404 | 0.0994 | 0.0337 | 0.0030 | 0.0190 | 0.0013 | 0.0089 | 0.0005 |
| 28 | 33.9488 | 0.2004 | 1.5466 | 0.0474 | 0.2011 | 0.0190 | 0.0704 | 0.0016 | 0.0123 | 0.0011 | 0.0081 | 0.0006 |

**Table S2.** Results for Mann-Whitney U test (MW) and bootstrap confidence interval on the median difference (BM) on the times at the *ops* site for varying forces (from 3 to 10 pN) in both AF and OF configurations.

| MW | -10 | -7 | -3 | 3 | 7 | 10 |
|---|---|---|---|---|---|---|
| -10 | — | $9.88 \times 10^{-1}$ | $8.30 \times 10^{-1}$ | $4.99 \times 10^{-1}$ | $6.61 \times 10^{-1}$ | $4.43 \times 10^{-1}$ |
| -7 | $9.88 \times 10^{-1}$ | — | $9.98 \times 10^{-1}$ | $2.59 \times 10^{-3}$ ** | $1.17 \times 10^{-1}$ | $3.27 \times 10^{-4}$ *** |
| -3 | $8.30 \times 10^{-1}$ | $9.98 \times 10^{-1}$ | — | $3.40 \times 10^{-4}$ *** | $3.92 \times 10^{-2}$ * | $1.29 \times 10^{-4}$ *** |
| 3 | $4.99 \times 10^{-1}$ | $2.59 \times 10^{-3}$ ** | $3.40 \times 10^{-4}$ *** | — | $3.55 \times 10^{-2}$ * | $7.99 \times 10^{-2}$ |
| 7 | $6.61 \times 10^{-1}$ | $1.17 \times 10^{-1}$ | $3.92 \times 10^{-2}$* | $3.55 \times 10^{-2}$ * | — | $2.93 \times 10^{-3}$ ** |
| 10 | $4.43 \times 10^{-1}$ | $3.27 \times 10^{-4}$ *** | $1.29 \times 10^{-4}$ *** | $7.99 \times 10^{-2}$ | $2.93 \times 10^{-3}$ ** | — |

| BM | -10 | -7 | -3 | 3 | 7 | 10 |
|---|---|---|---|---|---|---|
| -10 | — | $6.39 \times 10^{-1}$ | $5.34 \times 10^{-1}$ | $5.16 \times 10^{-1}$ | $5.23 \times 10^{-1}$ | $5.20 \times 10^{-1}$ |
| -7 | $6.39 \times 10^{-1}$ | — | $8.48 \times 10^{-1}$ | $7.42 \times 10^{-2}$ | $3.17 \times 10^{-1}$ | $4.34 \times 10^{-3}$** |
| -3 | $5.34 \times 10^{-1}$ | $8.48 \times 10^{-1}$ | — | $1.12 \times 10^{-3}$** | $5.54 \times 10^{-2}$ | $2.32 \times 10^{-3}$** |
| 3 | $5.16 \times 10^{-1}$ | $7.42 \times 10^{-2}$ | $1.12 \times 10^{-3}$** | — | $1.95 \times 10^{-1}$ | $6.01 \times 10^{-2}$ |
| 7 | $5.23 \times 10^{-1}$ | $3.17 \times 10^{-1}$ | $5.54 \times 10^{-2}$ | $1.95 \times 10^{-1}$ | — | $8.46 \times 10^{-3}$*** |
| 10 | $5.20 \times 10^{-1}$ | $4.34 \times 10^{-3}$** | $2.32 \times 10^{-3}$** | $6.01 \times 10^{-2}$ | $8.46 \times 10^{-3}$*** | — |

**Table S3.** Results for Mann-Whitney U test (MW) and bootstrap confidence interval on the median difference (BM) on the times at the *ops* site under 7 pN of force applied in OF configuration, in the presence of GreA (A), GreB (B), NusA (N), NusG (G) in isolation.

| MW | RNAP | A | B | N | G |
|---|---|---|---|---|---|
| **RNAP** | — | $8.09 \times 10^{-1}$ | $1.46 \times 10^{-2}$* | $2.73 \times 10^{-2}$* | $3.78 \times 10^{-1}$ |
| **A** | $8.09 \times 10^{-1}$ | — | $6.72 \times 10^{-4}$*** | $6.69 \times 10^{-3}$** | $2.04 \times 10^{-1}$ |
| **B** | $1.46 \times 10^{-2}$* | $6.72 \times 10^{-4}$*** | — | $7.44 \times 10^{-1}$ | $5.74 \times 10^{-5}$*** |
| **N** | $2.73 \times 10^{-2}$* | $6.69 \times 10^{-3}$** | $7.44 \times 10^{-1}$ | — | $5.09 \times 10^{-4}$*** |
| **G** | $3.78 \times 10^{-1}$ | $2.04 \times 10^{-1}$ | $5.74 \times 10^{-5}$*** | $5.09 \times 10^{-4}$*** | — |

| BM | RNAP | A | B | N | G |
|---|---|---|---|---|---|
| **RNAP** | — | $8.99 \times 10^{-01}$ | $8.60 \times 10^{-2}$ | $4.81 \times 10^{-2}$* | $5.95 \times 10^{-1}$ |
| **A** | $8.99 \times 10^{-01}$ | — | $1.20 \times 10^{-4}$*** | $1.23 \times 10^{-2}$* | $3.76 \times 10^{-1}$ |
| **B** | $8.60 \times 10^{-2}$ | $1.20 \times 10^{-4}$*** | — | $4.04 \times 10^{-1}$ | $8.00 \times 10^{-5}$*** |
| **N** | $4.81 \times 10^{-2}$* | $1.23 \times 10^{-2}$* | $4.04 \times 10^{-1}$ | — | $5.20 \times 10^{-3}$** |
| **G** | $5.95 \times 10^{-1}$ | $3.76 \times 10^{-1}$ | $8.00 \times 10^{-5}$*** | $5.20 \times 10^{-3}$** | — |

**Table S4.** Results for Mann-Whitney U test (MW) and bootstrap confidence interval on the median difference (BM) on the times at the *ops* site under 7 pN of force applied in OF configuration, in the presence of NusA (N), in combination with GreA (A) and/or NusG (G), or the latter two together.

| MW | N | NA | NG | GA | GNA |
|---|---|---|---|---|---|
| **N** | — | $8.58 \times 10^{-1}$ | $5.99 \times 10^{-1}$ | $2.26 \times 10^{-5}$*** | $3.53 \times 10^{-1}$ |
| **NA** | $8.58 \times 10^{-1}$ | — | $5.48 \times 10^{-1}$ | $1.03 \times 10^{-8}$*** | $1.95 \times 10^{-1}$ |
| **NG** | $5.99 \times 10^{-1}$ | $5.48 \times 10^{-1}$ | — | $3.21 \times 10^{-8}$*** | $6.75 \times 10^{-1}$ |
| **GA** | $2.26 \times 10^{-5}$*** | $1.03 \times 10^{-8}$*** | $3.21 \times 10^{-8}$*** | — | $1.74 \times 10^{-6}$*** |
| **GNA** | $3.53 \times 10^{-1}$ | $1.95 \times 10^{-1}$ | $6.75 \times 10^{-1}$ | $1.74 \times 10^{-6}$*** | — |

| BM | N | NA | NG | GA | GNA |
|---|---|---|---|---|---|
| **N** | — | $7.27 \times 10^{-1}$ | $4.73 \times 10^{-1}$ | $2.00 \times 10^{-5}$*** | $3.24 \times 10^{-1}$ |
| **NA** | $7.27 \times 10^{-1}$ | — | $2.49 \times 10^{-1}$ | $2.00 \times 10^{-5}$*** | $3.24 \times 10^{-1}$ |
| **NG** | $4.73 \times 10^{-1}$ | $2.49 \times 10^{-1}$ | — | $< 1 \times 10^{-5}$*** | $9.30 \times 10^{-1}$ |
| **GA** | $2.00 \times 10^{-5}$*** | $2.00 \times 10^{-5}$*** | $< 1 \times 10^{-5}$*** | — | $< 1 \times 10^{-5}$*** |
| **GNA** | $3.24 \times 10^{-1}$ | $3.24 \times 10^{-1}$ | $9.30 \times 10^{-1}$ | $< 1 \times 10^{-5}$*** | — |

**Table S5.** Results for Mann-Whitney U test (MW) and bootstrap confidence interval on the median difference (BM) on the times at the *ops* site under 7 pN of force applied in OF configuration, in the increasing concentrations (0 – 1 μM) of RfaH (H).

| **MW** | **0** | **0.005** | **0.05** | **0.25** | **1** |
|---|---|---|---|---|---|
| **0** | — | $5.91 \times 10^{-1}$ | $4.89 \times 10^{-1}$ | $3.88 \times 10^{-1}$ | $4.56 \times 10^{-2}$* |
| **0.005** | $5.91 \times 10^{-1}$ | — | $8.22 \times 10^{-1}$ | $2.71 \times 10^{-1}$ | $6.65 \times 10^{-2}$ |
| **0.05** | $4.89 \times 10^{-1}$ | $8.22 \times 10^{-1}$ | — | $2.17 \times 10^{-1}$ | $3.71 \times 10^{-2}$* |
| **0.25** | $3.88 \times 10^{-1}$ | $2.71 \times 10^{-1}$ | $2.17 \times 10^{-1}$ | — | $4.34 \times 10^{-1}$ |
| **1** | $4.56 \times 10^{-2}$* | $6.65 \times 10^{-2}$ | $3.71 \times 10^{-2}$* | $4.34 \times 10^{-1}$ | — |

| **BM** | **0** | **0.005** | **0.05** | **0.25** | **1** |
|---|---|---|---|---|---|
| **0** | — | $4.56 \times 10^{-1}$ | $4.41 \times 10^{-1}$ | $4.81 \times 10^{-1}$ | $6.30 \times 10^{-2}$ |
| **0.005** | $4.56 \times 10^{-1}$ | — | $9.79 \times 10^{-1}$ | $2.06 \times 10^{-1}$ | $1.75 \times 10^{-2}$* |
| **0.05** | $4.41 \times 10^{-1}$ | $9.79 \times 10^{-1}$ | — | $2.11 \times 10^{-1}$ | $2.24 \times 10^{-2}$* |
| **0.25** | $4.81 \times 10^{-1}$ | $2.06 \times 10^{-1}$ | $2.11 \times 10^{-1}$ | — | $2.31 \times 10^{-1}$ |
| **1** | $6.30 \times 10^{-2}$ | $1.75 \times 10^{-2}$* | $2.24 \times 10^{-2}$* | $2.31 \times 10^{-1}$ | — |

**Table S6.** Results for Mann-Whitney U test (MW) and bootstrap confidence interval on the median difference (BM) on the times at the *ops* site under 7 pN of force applied in OF configuration, in the increasing concentrations (0 – 1 μM) of RfaH (H).

| MW | -10 | -7 | -3 | 3 | 7 | 10 |
|---|---|---|---|---|---|---|
| **-10** | — | $5.42 \times 10^{-1}$ | $9.08 \times 10^{-1}$ | $9.85 \times 10^{-1}$ | $7.26 \times 10^{-1}$ | $2.09 \times 10^{-1}$ |
| **-7** | $5.42 \times 10^{-1}$ | — | $2.80 \times 10^{-2}$ * | $6.15 \times 10^{-1}$ | $2.56 \times 10^{-1}$ | $6.67 \times 10^{-3}$ ** |
| **-3** | $9.08 \times 10^{-1}$ | $2.80 \times 10^{-2}$ * | — | $3.47 \times 10^{-1}$ | $3.05 \times 10^{-1}$ | $4.43 \times 10^{-4}$ *** |
| **3** | $9.85 \times 10^{-1}$ | $6.15 \times 10^{-1}$ | $3.47 \times 10^{-1}$ | — | $9.85 \times 10^{-1}$ | $6.39 \times 10^{-3}$ ** |
| **7** | $7.26 \times 10^{-1}$ | $2.56 \times 10^{-1}$ | $3.05 \times 10^{-1}$ | $9.85 \times 10^{-1}$ | — | $1.59 \times 10^{-3}$ ** |
| **10** | $2.09 \times 10^{-1}$ | $6.67 \times 10^{-3}$ ** | $4.43 \times 10^{-4}$ *** | $6.39 \times 10^{-3}$ ** | $1.59 \times 10^{-3}$ ** | — |

| BM | -10 | -7 | -3 | 3 | 7 | 10 |
|---|---|---|---|---|---|---|
| **-10** | — | $4.45 \times 10^{-2}$ * | $5.60 \times 10^{-1}$ | $8.96 \times 10^{-1}$ | $3.72 \times 10^{-1}$ | $6.64 \times 10^{-3}$ ** |
| **-7** | $4.45 \times 10^{-2}$ * | — | $8.98 \times 10^{-2}$ | $2.47 \times 10^{-1}$ | $1.10 \times 10^{-1}$ | $2.09 \times 10^{-1}$ |
| **-3** | $5.60 \times 10^{-1}$ | $8.98 \times 10^{-2}$ | — | $6.75 \times 10^{-1}$ | $7.94 \times 10^{-1}$ | $8.78 \times 10^{-3}$** |
| **3** | $8.96 \times 10^{-1}$ | $2.47 \times 10^{-1}$ | $6.75 \times 10^{-1}$ | — | $5.32 \times 10^{-1}$ | $7.22 \times 10^{-2}$ |
| **7** | $3.72 \times 10^{-1}$ | $1.10 \times 10^{-1}$ | $7.94 \times 10^{-1}$ | $5.32 \times 10^{-1}$ | — | $1.03 \times 10^{-2}$ * |
| **10** | $6.64 \times 10^{-3}$ ** | $2.09 \times 10^{-1}$ | $8.78 \times 10^{-3}$** | $7.22 \times 10^{-2}$ | $1.03 \times 10^{-2}$* | — |

**Table S7.** Results for Mann-Whitney U test (MW) and bootstrap confidence interval on the median difference (BM) on the times at the *ops* site under 7 pN of force applied in OF configuration, in the presence of 1 µM of RfaH (H), with either 2 µM GreA (A), 2 µM GreB (B) or 10 nM nbRNAP.

| MW | H | HA | HB | H nbRNAP |
|---|---|---|---|---|
| **H** | — | $6.65 \times 10^{-4}$*** | $4.74 \times 10^{-3}$** | $1.20 \times 10^{-1}$ |
| **HA** | $6.65 \times 10^{-4}$*** | — | $6.71 \times 10^{-2}$ | $6.41 \times 10^{-2}$ |
| **HB** | $4.74 \times 10^{-3}$** | $6.71 \times 10^{-2}$ | — | $6.20 \times 10^{-1}$ |
| **H nbRNAP** | $1.20 \times 10^{-1}$ | $6.41 \times 10^{-2}$ | $6.20 \times 10^{-1}$ | — |

| BM | H | HA | HB | H nbRNAP |
|---|---|---|---|---|
| **H** | — | $6.80 \times 10^{-4}$*** | $2.83 \times 10^{-2}$* | $1.33 \times 10^{-1}$ |
| **HA** | $6.80 \times 10^{-4}$*** | — | $3.53 \times 10^{-2}$* | $6.62 \times 10^{-2}$ |
| **HB** | $2.83 \times 10^{-2}$* | $3.53 \times 10^{-2}$* | — | $7.00 \times 10^{-1}$ |
| **H nbRNAP** | $1.33 \times 10^{-1}$ | $6.62 \times 10^{-2}$ | $7.00 \times 10^{-1}$ | — |

**Table S8.** Results for Mann-Whitney U test (MW) and bootstrap confidence interval on the median difference (BM) on the times at the *ops* site under 7 pN of force applied in OF configuration, in the presence of NusA (N), RfaH (H) or their combination (HN).

| MW | N | H | HN |
|---|---|---|---|
| N | — | $2.88 \times 10^{-1}$ | $3.74 \times 10^{-1}$ |
| H | $2.88 \times 10^{-1}$ | — | $5.86 \times 10^{-1}$ |
| HN | $3.74 \times 10^{-1}$ | $5.86 \times 10^{-1}$ | — |

| BM | N | H | HN |
|---|---|---|---|
| N | — | $2.90 \times 10^{-1}$ | $4.40 \times 10^{-1}$ |
| H | $2.90 \times 10^{-1}$ | — | $8.13 \times 10^{-1}$ |
| HN | $4.40 \times 10^{-1}$ | $8.13 \times 10^{-1}$ | — |

**Table S9.** Results for Mann-Whitney U test (MW) and bootstrap confidence interval on the median difference (BM) on the times at the *ops* site under 7 pN of force applied in OF configuration, in the presence of NusG and GreA without (GA) or with NusA (GNA), or RfaH and GreA without (HA) or with NusA (HNA).

| MW | GA | GNA | HA | HNA |
|---|---|---|---|---|
| **GA** | — | $1.74 \times 10^{-6}$*** | $1.05 \times 10^{-5}$*** | $3.05 \times 10^{-14}$*** |
| **GNA** | $1.74 \times 10^{-6}$*** | — | $7.43 \times 10^{-2}$ | $1.13 \times 10^{-3}$** |
| **HA** | $1.05 \times 10^{-5}$*** | $7.43 \times 10^{-2}$ | — | $6.17 \times 10^{-1}$ |
| **HNA** | $3.05 \times 10^{-14}$*** | $1.13 \times 10^{-3}$** | $6.17 \times 10^{-1}$ | — |

| BM | GA | GNA | HA | HNA |
|---|---|---|---|---|
| **GA** | — | $2.00 \times 10^{-5}$*** | $3.80 \times 10^{-4}$*** | $< 1 \times 10^{-5}$*** |
| **GNA** | $2.00 \times 10^{-5}$*** | — | $1.70 \times 10^{-1}$ | $3.67 \times 10^{-2}$* |
| **HA** | $3.80 \times 10^{-4}$*** | $1.70 \times 10^{-1}$ | — | $3.56 \times 10^{-1}$ |
| **HNA** | $< 1 \times 10^{-5}$*** | $3.67 \times 10^{-2}$* | $3.56 \times 10^{-1}$ | — |

**Table S10.** List of primers used to prepare dsDNA constructs.

| Primer label | Sequence (5'-3') |
|---|---|
| fw AF handle | AAAAAAGCTTGGAACCAAAGGATATTCAGACGCG |
| rv AF handle | AAAAGGTCTCATTGCGGATCCCGTGATGACCTC |
| fw OF handle | AAAACCTAAGAGACCGGAACCAAAGGATATTCAGACG |
| rv OF handle | AAAAGGATCCCGTGATGACCTCATTA |
| fw AF stem | AAAATAGGAGAGACCGATAACGCAGGAAAGAACATGTG |
| rv AF stem | GTGGCCTTCTTCATCCAAGT |
| fw OF stem | CAGTGAATTCGAGCTCGGTA |
| rv OF stem | AATTGGTCTCTGCAAGGTCTTTCTTCGCCTGTTTG |

**Table S11.** Summary of the constrains for the parameters used to fit the dwell times distributions.

| **Parameter** | **Minimum value** | **Maximum value** |
|---|---|---|
| $k_{el}$ | 20.0 | 70.0 |
| $k_1$ | 1.0 | 5.0 |
| $k_2$ | 0.2 | 2.0 |
| $P_1$ | 0.1 | 0.5 |
| $P_2$ | 0.01 | 0.3 |
| $P_{pl}$ | 0.0 | 0.2 |

**Table S12.** Strains, genotype and source of the plasmids for *in vivo* assays.

| Strains | Genotype | Source |
|---|---|---|
| IA960 | *E. coli* MG1655 *argE*::Tn10 | This work |
| IA1016 | *E. coli* MG1655 *F' lacZY*::Tn9 | Natacha Ruiz |
| IA1017 | IA1016 *rfaH::*Kan$^R$ | This work |

**Sequence of pIA1437 template**

CGCGCGTTTCGGTGATGACGGTGAAAACCTCTGACACATGCAGCTCCCGGAGACGGTCACAGCTTGTCTGTAA
GCGGATGCCGGGAGCAGACAAGCCCGTCAGGGCGCGTCAGCGGGTGTTGGCGGGTGTCGGGGCTGGCTTA
ACTATGCGGCATCAGAGCAGATTGTACTGAGAGTGCACCATAAAATTGTAAACGTTAATATTTTGTTAAAATTCGCGT
TAAATTTTTGTTAAATCAGCTCATTTTTTAACCAATAGGCCGAAATCGGCAAAATCCCTTATAAATCAAAAGAATAGC
CCGAGATAGGGTTGAGTGTTGTTCCAGTTTGGAACAAGAGTCCACTATTAAAGAACGTGGACTCCAACGTCAAAG
GGCGAAAAACCGTCTATCAGGGCGATGGCCCACTACGTGAACCATCACCCAAATCAAGTTTTTTGGGGTCGAG
GTGCCGTAAAGCACTAAATCGGAACCCTAAAGGGAGCCCCCGATTTAGAGCTTGACGGGGAAAGCCGGCGAA
CGTGGCGAGAAAGGAAGGGAAGAAAGCGAAAGGAGCGGGCGCTAGGGCGCTGGCAAGTGTAGCGGTCACG
CTGCGCGTAACCACCACACCCGCCGCGCTTAATGCGCCGCTACAGGGCGCGTACTATGGTTGCTTTGACGTAT
GCGGTGTGAAATACCGCACAGATGCGTAAGGAGAAAATACCGCATCAGGCGCCATTCGCCATTCAGGCTGCG
CAACTGTTGGGAAGGGCGATCGGTGCGGGCCTCTTCGCTATTACGCCAGCTGGCGAAAGGGGGATGTGCTGC
AAGGCGATTAAGTTGGGTAACGCCAGGGTTTTCCCAGTCACGACGTTGTAAAACGACGGCCAGTGAATTCGAG
CTCGGTACCCGATCCAGATCCCGAACGCCTATCTTAAAGTTTAAACATAAAGACCAGACCTAAAGACCAGACCTA
AAGACACTACATAAAGACCAGACCTAAAGACGCCTTGTTGTTAGCCATAAAGTGATAACCTTTAATCATTGTCTTTAT
TAATACAACTCACTATAAGGAGAGACAACTTAAAGAGACTTAAAAGATTAATTTAAAATTTATCAAAAAGAGTAttgactT
AAAGTCTAACCTATAGgatactTACAGCCatcgagagggacacggggaaacaccaccaTCATCACCATCATCCTGACTAGAG
TGCTTGGCGAACCGGTGTTTGACGTCCAGGAATGTCAAATCCGTGGCGTGACCTATTCCGCACCGCTGCGCGT
TAAACTGCGTCTGGTGATCTATGAGCGCGAAGCGCCGGAAGGCACCGTAAAAGACATTAAAGAACAAGAAGTCT
ACATGGGCGAAATTCCGCTCATGACAGACAACGGTACCTTTGTTATCAACGGTACTGAGCGTGTTATCGTTTCCC
AGCTGCACCGTAGTCCGGGCGTCTTCTTTGACTCCGACAAAGGTAAAACCCACTCTTCGGGTAAAGTGCTGTAT
AACGCGCGTATCATCCCTTACCGTGGTTCCTGGCTGGACTTCGAATTCGATCCGAAGGACAACCTGTTCGTACCa
AtTGCGAAAGAAATGATACACTAGCACGTCAAAGTAAGTGCGTTATCAGTATTCAGGTAGCTGTTGAGCCTGGGGC
GGTAGCGTGCTTTTTTCTGCTTAACTTAACCAGACAATCACACAAAAGAGTCGCTAGTGGAAAAGCCATTTCGAAA
AATCCTGGTCATAAAGatgCGATATCATGGGGATATGTTATTAACTACTCCTGTCATCAGTACGCTCAAGCAGAATTA
TCCTGATGCAAAAATCGATATGCTGCTTTAaTCGACCGTCGCCGTAAACTGCCTGCGACCATCATTCTGCGCGCC
CTGAACTACACCACAGAGCAGATCCTCGACCTGTTtTTTGAAAAAGTTATCTTTGAAATCCGTGATAACAAGCTGCA
GATGGAACTGGTGCCGGAACGCCTGCGTGGTGAAACCGCATCTTTTGACATCGAAGCTAACGGTAAAGTGTACG
TAGAAAAAGGCCGCCGTATCACTGCGCGCCACATTCGCCAGCTGGAAAAAGACGACGTCAAACTGATCGAAGT
CCCGGTTGAGTACATCGCAGGTAAAGTGGTTGCTAAAGACTATATTGATGAGTCTACCGGCGAGCTGATCTGCGC
AGCGAACATGGAGCTGAGCCTGGATCTGCTGGCTAAGCTGAGCCAGTCTGGGAGCCTGGGGCGGTAGCGTG
CTTTTTTCTTCACCAACGATCTGGATCACGGCCCATATATCTCTGAAACCTTACGTGTCGACCCAACTAACGACCG
TCTGAGCGCACTGGTAGAAATCTACCGCATGATGCGCCCTGGCGAGCCGCCGACTCGTGAAGCAGCTGAAAG
CCTGTTCGAGAACCTGTTCTTCTCCGAAGACCGTTATGACTTGTCTGCGGTTGGTCGTATGAAGTTCAACCGTTCT
CTGCTGCGCGAAGAAATCGAAGGTTCCGGTATCCTGAGCAAAGACGACATCATTGATGTTATGAAAAAGCTCATC
GATATCCGTAACGGTAAAGGCGAAGTCGATGATATCGACCACCTCGGCAACCGTCGTATCCGTTCCGTTGGCGA
AATGGCGGAAAACCAGTTCCGCGTTGGCCTGGTACGTGTAGAGCGTGCGGTGAAAGAGCGTCTGTCTCTGGGC
GATCTGGATACCCTGATGCCACAGGATATGATCAACGCCAAGCCGATTTCCGCAGCAGTGAAAGAGTTCTTCGG
TTCCAGCCAGCTGTCTCAGTTTATGGACCAGAACAACCCGCTGTCTGAGATTACGCACAAACGTCGTATCTCCG
CACTCGGCCCAGGCGGTCTGACCCGTGAACGTGCAGGCTTCGAAGTTCGAGACGTACACCCGACTCACTACG
GTCGCGTATGTCCAATCGAAACCCCTGAAGGTCCGAACATCGGTCTGATCAACTCTCTGTCCGTGTACGCACAG
ACTAACGAATACGGCTTCCTTGAGACTCCGTATCGTAAAGTGACCGACGGTGTTGTAACTGACGAAATTCACTAC
CTGTCTGCTATCGAAGAAGGCAACTACGTTATCGCCCAGGCGAACTCCAACTTGGATGAAGAAGGCCACTTCGT
AGAAGACCTGGTAACTTGCCGTAGCAAAGGCGAATCCAGCTTGTTCAGCCGCGACCAGGTTGACTACATGGAC
GTATCCACCCAGCAGGTGGTATCCGTCGGTGCGTCCCTGATCCCGTTCCTGGAACACGATGACGCCAACCGT
GCATTGATGGGTGCGAACATGCAACGTCAGGCCGTTCCGACTCTGCGCGCTGATAAGCCGCTGGTTGGTACTG
GTATGGAACGTGCTGTTGCCGTTGACTCCGGTGTAACTGCGGTAGCTAAACGTGGTGGTGTCGTTCAGTACGTGG

ATGCTTCCCGTATCGTTATCAAAGTTAACGAAGACGAGATGTATCCGGGTGAAGCAGGTATCGACATCTACAACCT
GACCAAATACACCCGTTCTAACCAGAACACCTGTATCAACCAGATGCCGTGTGTGTCTCTGGGTGAACCGGTTG
AACGTGGCGACGTGCTGGCAGACGGTCCGTCCACCGACCTCGGTGAACTGGCGCTTGGTCAGAACATGCGC
GTAGCGTTCATGCCGTGGAATGGTTACAACTTCGAAGACTCCATCCTCGTATCCGAGCGTGTTGTTCAGGAAGAC
CGTTTCACCACCATCCACATTCAGGAACTGGCGTGTGTGTCCCGTGACACCAAGCTGGGTCCGGAAGAGATCA
CCGCTGACATCCCGAACGTGGGTGAAGCTGCGCTCTCCAAACTGGATGAATCCGGTATCGTTTACATTGGTGCG
GAAGTGACCGGTGGCGACATTCTGGTTGGTAAGGTAACGCCGAAAGGTGAAACTCAGCTGACCCCAGAAGAAA
AACTGCTGCGTGCGATCTTCGGTGAGAAAGCCTCTGACGTTAAAGACTCTTCTCTGCGCGTACCAAACGGTGTAT
CCGGTACGGTTATCGACGTTCAGGTCTTTACTCGCGATGGCGTAGAAAAAGACAAACGTGCGCTGGAAATCGAA
GAAATGCAGCTCAAACAGGCGAAGAAAGACCTGTCTGAAGAACTGCAGATCCTCGAAGCGGGTCTGTTCAGCC
GTATCCGTGCTGTGCTGGTAGCCGGTGGCGTTGAAGCTGAGAAGCTCGACAAACTGCCGCGCGATCGCTGGC
TGGAGCTGGGCCTGACAGACGAAGAGAAACAAAATCAGCTGGAACAGCTGGCTGAGCAGTATGACGAACTGAA
ACACGAGTTCGAGAAGAAACTCGAAGCGAAACGCCGCAAAATCACCCAGGGCGACGATCTGGCACCGGGCG
TGCTGAAGATTGTTAAGGTATATCTGGCGGTTAAACGCCGTATCCAGCCTGGTGACAAGATGGCAGGTCGTCACG
GTAACAAGGGTGTAATTTCTAAGATCAACCCGATCGAAGATATGCCTTACGATGAAAACGGTACGCCGGTAGACA
TCGTACTGAACCCGCTGGGCGTACCGTCTCGTATGAACATCGGTCAGATCCTCGAAACCCACCTGGGTATGGCT
GCGAAAGGTATCGGCGACAAGATCAACGCCATGCTGAAACAGCAGCAAGAAGTCGCGAAACTGCGCGAATTC
ATCCAGCGTGCGTACGATCTGGGCGCTGACGTTCGTCAGAAAGTTGACCTGAGTACCTTCAGCGATGAAGAAGT
TATGCGTCTGGCTGAAAACCTGCGCAAAGGTATGCCAATCGCAACGCCGGTGTTCGACGGTGCGAAAGAAGCA
GAAATTAAAGAGCTGCTGAAACTTGGCGACCTGCCGACTTCCGGTCAGATCCGCCTGTACGATGGTCGCACTG
GTGAACAGTTCGAGCGTCCGGTAACCGTTGGTTACATGTACATGCTGAAACTGAACCACCTGGTCGACGACAAG
ATGCACGCGCGTTCCACCGGTTCTTACAGCCTGGTTACTCAGCAGCCGCTGGGTGGTAAGGCACAGTTCGGTG
GTCAGCGTTTCGGGGAGATGGAAGTGTGGGCGCTGGAAGCATACGGCGCAGCATACACCCTGCAGGAAATGC
TCACCGTTAAGTCTGATGACGTGAACGGTCGTACCAAGATGTATAAAAACATCGTGGACGGCAACCATCAGATGG
AGCCGGGCATGCGAGAGTAGGGAACTGCCAGGCATCAAAGAAAACGAAAGGCACAGTCGAAAGACTGGGCC
TTTCGTTTTATCTGTTGTTTGTCGGTGAACGCTCTCCTGAGTAGGACAAATCCGCCGGGAGCGGATTTGAACGTTG
CGAAGCAACGGCCCGGAGGGTGGCGGGCAGGACGCCCGCCATAAACTGCCAGGCATCAAATTAAGCAGAA
GGCCATCCTGACGGATGGCCTTTTTGCGTTTCTACAAACTCTTCCTGTCGTCATATCTACAAGCCATCCCCCCAC
AGATACGGTAAACTAGCCTCGTTTTTGCATCAGGAAAGCAGAAGCTTGGCGTAATCATGGTCATAGCTGTTTCCTG
TGTGAAATTGTTATCCGCTCACAATTCCACACAACATACGAGCCGGAAGCATAAAGTGTAAAGCCTGGGGTGCCT
AATGAGTGAGCTAACTCACATTAATTGCGTTGCGCTCACTGCCCGCTTTCCAGTCGGGAAACCTGTCGTGCCAG
CTGCATTAATGAATCGGCCAACGCGCGGGGAGAGGCGGTTTGCGTATTGGGCGCTCTTCCGCTTCCTCGCTCA
CTGACTCGCTGCGCTCGGTCGTTCGGCTGCGGCGAGCGGTATCAGCTCACTCAAAGGCGGTAATACGGTTATC
CACAGAATCAGGGGATAACGCAGGAAAGAACATGTGAGCAAAAGGCCAGCAAAAGGCCAGGAACCGTAAAAA
GGCCGCGTTGCTGGCGTTTTTCCATAGGCTCCGCCCCCCTGACGAGCATCACAAAAATCGACGCTCAAGTCA
GAGGTGGCGAAACCCGACAGGACTATAAAGATACCAGGCGTTTCCCCCTGGAAGCTCCCTCGTGCGCTCTCC
TGTTCCGACCCTGCCGCTTACCGGATACCTGTCCGCCTTTCTCCCTTCGGGAAGCGTGGCGCTTTCTCATAGCT
CACGCTGTAGGTATCTCAGTTCGGTGTAGGTCGTTCGCTCCAAGCTGGGCTGTGTGCACGAACCCCCCGTTCA
GCCCGACCGCTGCGCCTTATCCGGTAACTATCGTCTTGAGTCCAACCCGGTAAGACACGACTTATCGCCACTG
GCAGCAGCCACTGGTAACAGGATTAGCAGAGCGAGGTATGTAGGCGGTGCTACAGAGTTCTTGAAGTGGTGGC
CTAACTACGGCTACACTAGAAGGACAGTATTTGGTATCTGCGCTCTGCTGAAGCCAGTTACCTTCGGAAAAAGAG
TTGGTAGCTCTTGATCCGGCAAACAAACCACCGCTGGTAGCGGTGGTTTTTTTGTTTGCAAGCAGCAGATTACGC
GCAGAAAAAAAGGATCTCAAGAAGATCCTTTGATCTTTTCTACGGGGTCTGACGCTCAGTGGAACGAAAACTCAC
GTTAAGGGATTTTGGTCATGAGATTATCAAAAAGGATCTTCACCTAGATCCTTTTAAATTAAAAATGAAGTTTTAAATC
AATCTAAAGTATATATGAGTAAACTTGGTCTGACAGTTACCAATGCTTAATCAGTGAGGCACCTATCTCAGCGATCTG
TCTATTTCGTTCATCCATAGTTGCCTGACTCCCCGTCGTGTAGATAACTACGATACGGGAGGGCTTACCATCTGGC
CCCAGTGCTGCAATGATACCGCGAGACCCACGCTCACCGGCTCCAGATTTATCAGCAATAAACCAGCCAGCC
GGAAGGGCCGAGCGCAGAAGTGGTCCTGCAACTTTATCCGCCTCCATCCAGTCTATTAATTGTTGCCGGGAAG
CTAGAGTAAGTAGTTCGCCAGTTAATAGTTTGCGCAACGTTGTTGCCATTGCTACAGGCATCGTGGTGTCACGCTC

GTCGTTTGGTATGGCTTCATTCAGCTCCGGTTCCCAACGATCAAGGCGAGTTACATGATCCCCCATGTTGTGCAA
AAAAGCGGTTAGCTCCTTCGGTCCTCCGATCGTTGTCAGAAGTAAGTTGGCCGCAGTGTTATCACTCATGGTTATG
GCAGCACTGCATAATTCTCTTACTGTCATGCCATCCGTAAGATGCTTTTCTGTGACTGGTGAGTACTCAACCAAGT
CATTCTGAGAATAGTGTATGCGGCGACCGAGTTGCTCTTGCCCGGCGTCAATACGGGATAATACCGCGCCACAT
AGCAGAACTTTAAAAGTGCTCATCATTGGAAAACGTTCTTCGGGGCGAAAACTCTCAAGGATCTTACCGCTGTTG
AGATCCAGTTCGATGTAACCCACTCGTGCACCCAACTGATCTTCAGCATCTTTTACTTTCACCAGCGTTTCTGGGT
GAGCAAAAACAGGAAGGCAAAATGCCGCAAAAAAGGGAATAAGGGCGACACGGAAATGTTGAATACTCATACT
CTTCCTTTTTCAATATTATTGAAGCATTTATCAGGGTTATTGTCTCATGAGCGGATACATATTTGAATGTATTTAGAAAAA
TAAACAAATAGGGGTTCCGCGCACATTTCCCCGAAAAGTGCCACCTGACGTCTAAGAAACCATTATTATCATGAC
ATTAACCTATAAAAATAGGCGTATCACGAGGCCCTTTCGTCT
